# Distinct chromatin scanning modes lead to targeting of compacted chromatin by pioneer factors FOXA1 and SOX2

**DOI:** 10.1101/2022.12.22.521655

**Authors:** Jonathan Lerner, Andrew Katznelson, Jingchao Zhang, Kenneth S. Zaret

**Affiliations:** Institute for Regenerative Medicine and Department of Cell and Developmental Biology, Perelman School of Medicine, University of Pennsylvania, Philadelphia, PA 19104-6058

**Author notes:** These authors contributed equally (Jonathan Lerner, Andrew Katznelson).

**Keywords:** chromatin, dynamics, pioneer factor, gene networks, reprogramming, development

## Abstract

Pioneer transcription factors, by interacting with nucleosomes, can scan silent, compact chromatin to target gene regulatory sequences, enabling cooperative binding events that modulate local chromatin structure and gene activity. However, pioneer factors do not target all of their cognate motifs and it is unclear whether different pioneers scan compact chromatin the same way. Surprisingly, combined approaches of genomics and single-molecule tracking show that to target DNase-resistant, low-histone turnover sites, pioneer factors can use opposite dynamics of chromatin scanning. FOXA1 uses low nucleoplasmic diffusion and stable chromatin interactions, whereas SOX2 uses high nucleoplasmic diffusion and transient interactions, respectively. Despite such differences, FOXA1 and SOX2 scan low-mobility, silent chromatin to similar extents, as mediated by protein domains outside of the respective DNA binding domains. By contrast, the non-pioneer HNF4A predominantly targets DNase-sensitive, nucleosome-depleted regions. We conclude that the targeting of compact chromatin sites by pioneer factors can be through distinct dynamic processes.

## INTRODUCTION

At the onset of cell fate transitions, genes that maintain the cell state of origin can become repressed, while genes required for the future cell state are activated. The rewiring of genetic networks can be driven by pioneer transcription factors, which bind to active and silent chromatin regions and recruit chromatin remodelers, coactivators, or corepressor complexes to elicit gene expression changes ^1–3^. Notably, pioneer factors enable zygotic genome activation in fruit flies ^4,5^, zebrafish ^6^, and mouse embryos ^7^. Pioneer transcription factors characteristically have DNA binding domains compatible with recognition of DNA motifs that may only be partially exposed on the nucleosome ^8–10^, which enables the targeting of silent chromatin with low levels of active or repressive histone marks ^11–13^. Whether nucleosome turnover and transiently free DNA in silent chromatin allows preferential targeting by pioneer factors is unclear, as is whether the factors, as a class, scan chromatin similarly or by different modalities.

Prior to binding to specific chromatin targets, pioneer and non-pioneer transcription factors perform an exploratory scanning of chromatin, alternating between nucleoplasmic diffusion and non-specific DNA and chromatin sampling ^14–16^. Fluorescence Recovery After Photobleaching (FRAP) showed that the pioneer factor FOXA1 presents lower nuclear dynamics than non-pioneers cMYC or NF-1, which was interpreted to indicate a slow, lateral scanning across nucleosome-dense chromatin ^17^. The development of Single-Molecule Tracking (SMT) has allowed a direct assessment of chromatin scanning by transcription factors, revealing spatiotemporal parameters of nucleoplasmic diffusion and residence times ^14^. While the Drosophila pioneer factor GAGA presents residence times higher than most transcription factors ^5^, most pioneer transcription factors, including FOXA1, presents residence times on chromatin like non-pioneers, in the range of 10-20 seconds ^14,18–20^.

Using SMT ^14,21^, we developed a two-parameter method for analyzing chromatin motions, as initially defined in other studies ^22–24^, where the motions can be functionally classified into a range of high to low mobility chromatin domains ^19^. The low-mobility domains are bound by heterochromatin-associated proteins such as HP1a, Lamin A, and H3K9me3 histone methyltransferases ^19^. Among nine transcription factors tested in a mouse hepatic cell line, nucleosome-binding pioneer factors FOXA1 and SOX2 presented the strongest ability to bind and scan low-mobility chromatin, while the weak-nucleosome binding transcription factor HNF4A was primarily found in high-mobility chromatin ^19^. Using a different methodological approach, a recent SMT-based study found that the glucocorticoid receptor binds to regions of low and high confinement after induction in a mouse hepatocarcinoma cell line ^25^.

The SMT analysis of ectopically expressed FOXA1 in hepatic cells ^19^ could have been biased by conditioning of the chromatin environment due to the endogenous expression of FOXA1 and co-acting factors ^26–28^. The cellular context in which transcription factors are studied can influence their activity ^29^. Yet in an ectopic expression context, evidence exists for transient or low-level “sampling” of alternate sites not stably bound in one cell type ^11^. The extent to which such sampling may depend on specific DNA site recognition is unclear.

Here we use orthogonal approaches combining ChIP-seq, CUT&RUN, and SMT of pioneer factors FOXA1 and SOX2 and the non-pioneer HNF4A to address three main questions. First, while pioneer factors may target compact chromatin that is DNase-resistant, do they target sites of low or high histone turnover? Models where transcription factors obligatorily target sites of free DNA would be expected to occur at sites of higher histone turnover. If different pioneer factors target comparable fractions of DNase-resistant sites, do they do so by similar or distinct chromatin scanning mechanisms? Is nucleosome binding by the DNA binding domain of pioneer factors sufficient to enable closed chromatin scanning and targeting? The results reveal an unexpected diversity in molecular dynamics by which pioneer factors scan and target closed chromatin domains, providing insights into different ways that cell fates can be reprogrammed.

## RESULTS

### Targeting of low nucleosome turnover sites by FOXA1 and SOX2, but not HNF4A

To assess how pioneer versus non-pioneer transcription factors target a novel chromatin landscape, we ectopically introduced FOXA1-HALO, HNF4A-HALO or SOX2-V5 in primary human fibroblasts for 48 hours using a lentiviral expression system (Figure 1A and Supplemental Figure 1A). As these factors are not endogenously expressed in fibroblasts, evaluating the initial ectopic binding events assesses their respective capacity to “pioneer” distinct chromatin states. By expressing the factors de novo in each experiment, we eliminate possible “priming” artifacts due to low basal expression, as can be seen with inducible vectors. We also carefully titrated expression levels to be comparable among the factors (Supplemental Figure 1A).

**Figure 1:**
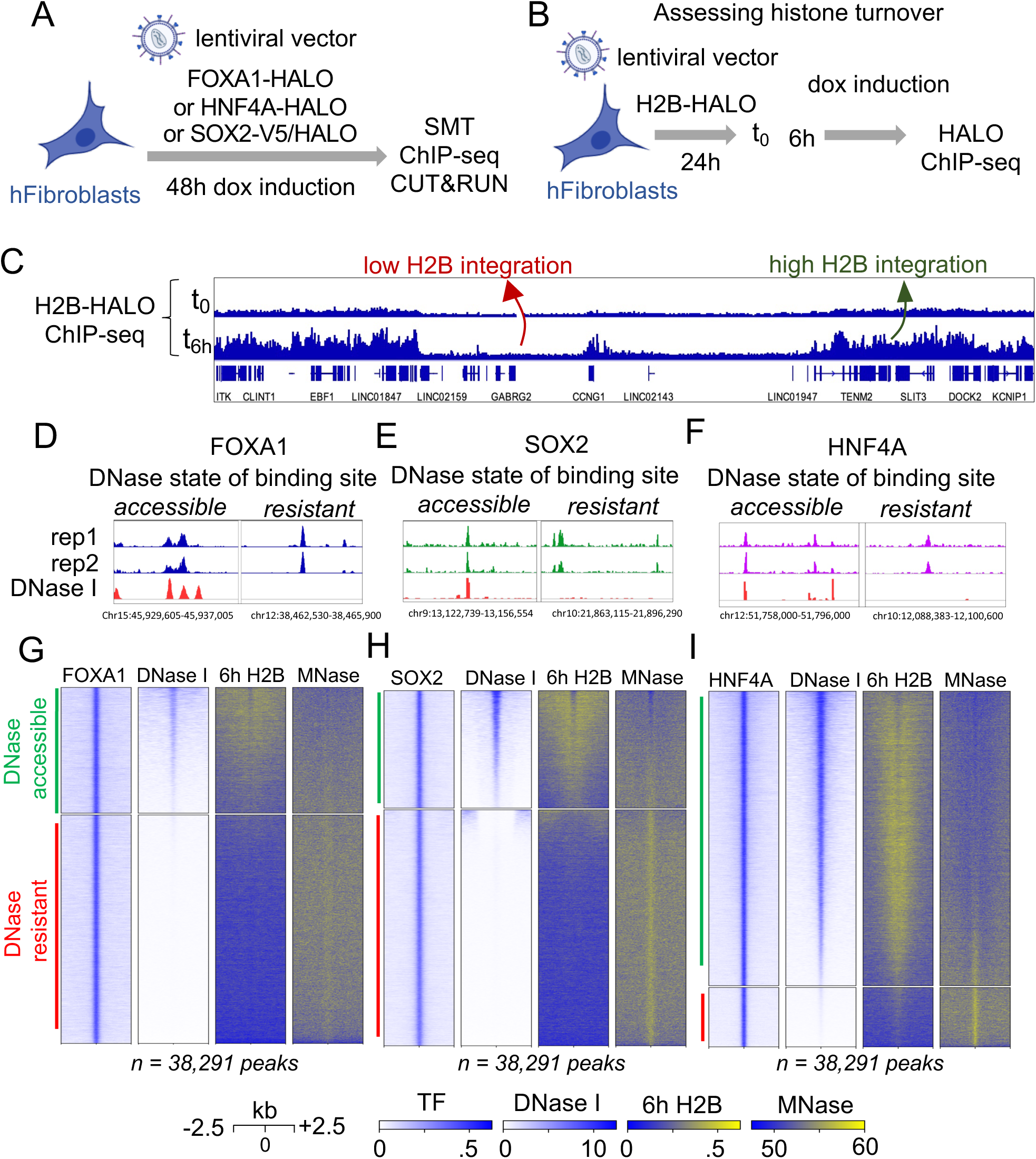
Ectopically expressed pioneer factors FOXA1 and SOX2 predominantly bind to silent, low histone integration chromatin. A: Experimental strategy to dissect chromatin interactions of transcription factors. FOXA1-HALO, SOX2-V5/HALO, and HNF4A-HALO were ectopically expressed in human fibroblasts with lentiviral vectors for 48 hours, followed by ChIP-seq (FOXA1, HNF4A), CUT&RUN (SOX2) and SMT (FOXA1, HNF4A, and SOX2). B: Experimental strategy to annotate regions of high and low histone turnover. After 24 hours of infection with lentiviral vectors, H2B-HALO expression is induced with doxycycline for 6 hours, and ChIP-seq is performed against the HALO tag. C: Browser track views show regions of low versus high H2B-HALO integration. D-F: Examples of ChIP-seq (FOXA1, HNF4A) or CUT&RUN (SOX2) peaks stratified by overlap with DNase I sensitive (accessible) or resistant chromatin. G-I: Heatmaps displaying the pre-existing chromatin features in human fibroblasts, centered on ectopic FOXA1 (G), HNF4A (H) and SOX2 (I) target sites. FOXA1 (G) and SOX2 (I) peaks were randomly down-sampled to the number of HNF4A peaks (n=38,291).

We first assessed whether the factors target silent DNA sites on labile or stable nucleosomes, and whether high histone dynamics is a precondition for pioneer factor binding, even in closed chromatin. To this end, we pulsed histone H2B-HALO expression for 6 hours in human fibroblasts and performed ChIP-seq against the HALO tag (Figure 1B), with enrichment marking regions that have integrated histones during the time period (Figure 1C). Next, we mapped nucleosome positions by digesting chromatin with high concentrations of micrococcal nuclease (MNase), isolating mononucleosome-sized fragments, and sequencing the underlying DNA (Supplemental Figure 1B and 1C) ^26^. The concordance of histone turnover and nucleosome positioning annotations, along with integration of DNase-seq data ^30^, allowed us to define the pre-existing nucleosome and chromatin accessibility states prior to an ectopic transcription factor binding event (Supplemental Figure 1D and E). Using ENCODE annotations of candidate cis regulatory elements (cCREs) ^31^ and in agreement with prior studies ^32,33^, we find that the centers of active promoter cCREs in fibroblasts are nucleosome free, with immediately flanking domains showing high histone turnover and one or two positioned nucleosomes, while inactive cCREs at enhancers are in low-turnover nucleosome domains (Supplemental Figure 1D-E).

To assess chromatin targeting by the three transcription factors, we performed ChIP-seq for FOXA1-HALO and HNF4A-HALO, and CUT&RUN ^34^ for SOX2-V5 (Supplemental Figure 2A). We assessed the reproducibility of our ChIP-seq and CUT&RUN experiments by mapping FOXA1-HALO and SOX2-V5 signals over FOXA2 and SOX2 peaks profiled in two previous studies, after ectopic expression in human fibroblasts ^11,13^, and observe a high concordance between datasets (Supplemental Figure 2B and C). Here we analyzed the factors in parallel, in fibroblasts.

To classify binding events in open versus closed chromatin, we stratified the peak sets by overlap with DNase-I hypersensitivity or insensitivity (Figure 1D-F). For a side-by-side comparison between the three transcription factors, we randomly down-sampled FOXA1 and SOX2 peaks to the number of HNF4a binding sites (Figure 1G-I and Supplemental Figure 2D, E). We find that the majority of FOXA1 (64.5%, Figure 1G and Supplemental Figure 2D) and SOX2 (67%, Figure 1H and Supplemental Figure 2E) targeting events are in DNase I-resistant chromatin, before or after down-sampling. Sites and domains targeted in DNase-resistant chromatin displayed MNase-resistant signals (Figure 1G-I), confirming the targeting of pioneer factors to nucleosomal DNA. Interestingly, the MNase signals directly underlying FOXA1 binding events in DNase-resistant chromatin (Figure 1G) appeared fuzzier than for SOX2 (Figure 1H and I), which may reflect a different dependence of a positioned nucleosome for targeting by the factors. By contrast, a much smaller fraction of comparably expressed (Supplemental Figure 1A) HNF4a targeted sites (16.7%) fall in DNase-resistant chromatin (Figure 1I).

The targeted sites in DNase-sensitive chromatin had high histone turnover and low MNase-resistant signals, reflecting the dynamic nature of open chromatin states (Figure 1G-The fewer DNase resistant sites targeted by HNF4A show histone turnover (Figure 1I, Supplemental Figure 2F, blue line). Thus, by the static assays of either ChIP or CUT&RUN, the pioneer factors FOXA1 and SOX2 preferentially target silent, low dynamic chromatin states, while HNF4A mainly targets open sites with more dynamic chromatin.

### Distinct chromatin scanning properties can lead to compact chromatin occupancy

We next addressed whether the common extent of closed chromatin targeting by FOXA1 and SOX2 (64-67%) occur via similar or different chromatin scanning dynamics (Figure 2A). We employed single molecule tracking (SMT) ^21, 35–37^ ^14,38^ of individual molecules of FOXA1-HALO, SOX2-HALO, and HNF4A-HALO after ectopic expression in human fibroblasts (Supplemental Figure 1A and 3A).

**Figure 2:**
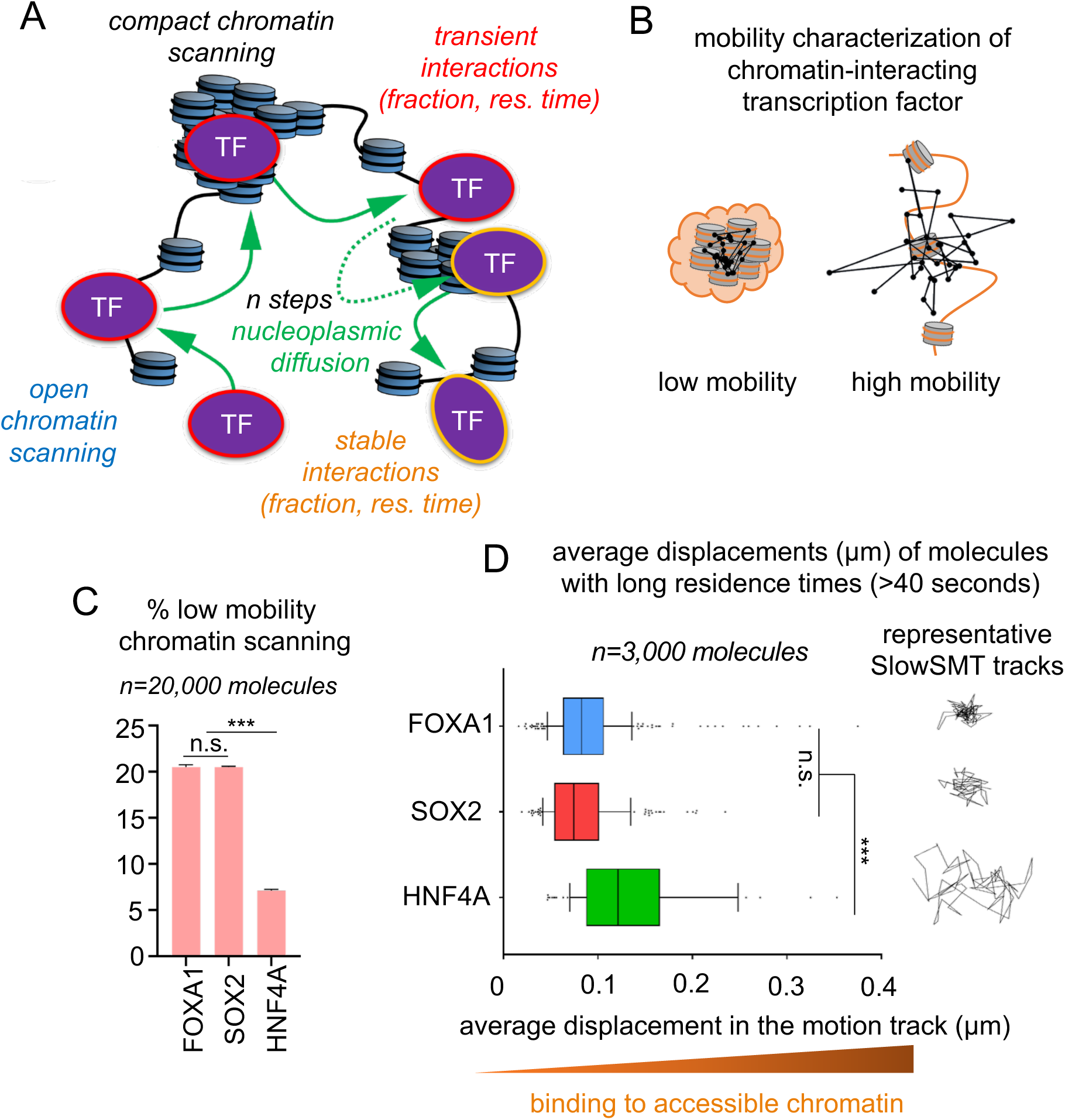
FOXA1 and SOX2 display enhanced scanning interactions with low-mobility chromatin. A: Process of chromatin scanning by transcription factors: Transcription factors alternate between nucleoplasmic diffusion (green) and chromatin interactions. Chromatin interactions occur in open (blue) or compact (black) chromatin, with transient or stable residence times. B: In living cells, pioneer transcription factors interact with low (left) and high (right) mobility chromatin. Low mobility chromatin is characteristically bound by heterochromatin regulators, which suggests a compact state. Non-pioneers are found interacting primarily with high mobility chromatin. C: SMT measurement of scanning of low mobility chromatin by FOXA1-HALO, SOX2-HALO and HNF4A-HALO. *** indicates p<0.0001, n.s. non-significant differences (p>0.05) as determined by one-way ANOVA, see Table S1). D: Average displacements of SlowSMT motion tracks of FOXA1-HALO (blue) SOX2-HALO (red) and HNF4A-HALO (green) for molecules with long residence times. Long residence times are defined as above 40 seconds, corresponding to the longest binding events that can be measured by SlowSMT, representing the 95^th^ percentile of histone H2B residence time distribution (see Supplemental Figure 4A).Representative motion tracks are shown for each transcription factors. Increased displacements reflect binding to more open chromatin. *** indicates p<0.0001, n.s. non-significant differences (p>0.05) as determined by one-way ANOVA, see Table S1).

Using FastSMT ^14^, we first measured the logarithmic distribution of the diffusion coefficients (in μm2 /s) of the motion tracks of the transcription factors. FastSMT infers and quantifies whether molecules are interacting with chromatin or diffusing in the nucleoplasm ^14,39^, as assessed by their overlapping with profiles for histone H2B or dCas9 expressed without a guide RNA, respectively (Supplemental Figure 3B-C) ^19^. As seen previously in hepatic cells ^17,19^, in human fibroblasts we observed that FOXA1 presents low levels of nucleoplasmic diffusion and high levels of chromatin interactions, whereas HNF4A and SOX2 present higher levels of nucleoplasmic diffusion and lower levels of chromatin interactions (Supplemental Figure 3D). However, applying a two-parameter mobility analysis that assesses the underlying chromatin dynamics by comparing the average displacement versus the radius of confinement of a single transcription factor molecule (Figure 2B), ^40^, we found that chromatin-bound FOXA1 and SOX2 engage in comparably higher fractions of low-mobility chromatin scanning (~20%) versus HNF4A (~7%) (Figure 2C, and primary data in Supplemental Figure 3E). Thus, both the ChIP-seq and CUT&RUN data for the two pioneer factors reveal a correspondence between targeting of DNase-I resistant chromatin with scanning of low-mobility chromatin.

We next aimed to assess the chromatin dynamics underlying highly stable binding events of FOXA1, SOX2, and HNF4A. We used SlowSMT ^39^ to track stably bound molecules of FOXA1, HNF4A, and SOX2 and measure their residence times (Supplemental Figure 4A), using a photobleaching correction based on the distribution of histone H2B residence times ^18,41^ (see STAR Methods). To infer how chromatin dynamics influences the binding stability of FOXA1, SOX2, and HNF4A, we associated the dwell times of each SlowSMT motion track with its corresponding average displacement in the track, since shorter displacements reflect binding to low mobility chromatin states (see STAR Methods). We then defined a pool of transcription factor molecules showing high binding stability, above the 99^th^ percentile of Histone H2B residence times (40 seconds, Supplemental Figure 4A).

We observed that the SlowSMT motion tracks of high binding-stability molecules of HNF4A presented significantly higher displacement than comparably expressed FOXA1 and SOX2 (Figure 2D, Supplemental Figure 1), reflecting how the latter pioneer factors establish stable interactions with low mobility chromatin states. Even HNF4A molecules with lower residence times displayed a significant increase in displacements compared to the two pioneer transcription factors (Supplemental Figure 4B). Of note, while the measurement of displacements reveals binding to high and low mobility chromatin, FOXA1 displays more longer-lived binding events (Supplemental Figure 4A, C-E) compared to SOX2 and HNF4A, correlating with a higher rate of chromatin interactions (Supplemental Figure 3D) and reflecting a mode of chromatin scanning based on a greater number of stable interactions. By contrast, HNF4A displays fast scanning dynamics and a reduced capacity to establish stable interactions in low-mobility chromatin.

Taken together, the results show that pioneers FOXA1 and SOX2 employ distinct dynamics to scan low-mobility chromatin. FOXA1 has a slow-scanning behavior, with low nucleoplasmic diffusion and stable chromatin interactions, while SOX2 has a fast-scanning behavior, with high nucleoplasmic diffusion and more transient chromatin interactions. Thus, our observations indicate that a pioneer factor can use one modality or another to target silent, DNase I resistant, compact chromatin sites.

### Integration of single molecule data to simulate transcription factor scanning

We developed a discrete-step simulation using parameters derived from FOXA1, HNF4A, and SOX2 SMT data to visualize the chromatin scanning process (Figure 3A). We integrated: a) the percent of molecules in nucleoplasmic diffusion (P_diffusion_) or chromatin interactions (Figure 3A, P_diffusion_ and P_interaction_ and Supplemental Figure 3D); b) the average diffusing distance (Supplemental Figure 4F and G and see STAR Methods); and c) engagement with low-mobility chromatin (Figure 2C), to generate spatial trajectories of single molecules traveling to 1,000 nonspecific chromatin sites (Figure 3B-D). To define the total time spent by the molecule scanning 1,000 chromatin sites, we summed 1,000 randomly selected dwell times from the distribution of residence times for FOXA1, SOX2, and HNF4A (Supplemental Figure 4A). To estimate the time spent in diffusion, we used the average duration of sorted motion tracks of cells performing nucleoplasmic diffusion (Supplemental Figure 4H, I and see STAR Methods). We did not make assumptions relating dwell times to interaction states (e.g. specific versus nonspecific binding), notably because it is unclear whether SMT truly captures residence times corresponding to specific DNA binding ^18,42–44^. Our simulations allow us to integrate multiple data modalities from SMT measurements to model and visualize chromatin scanning activities between different transcription factors.

**Figure 3:**
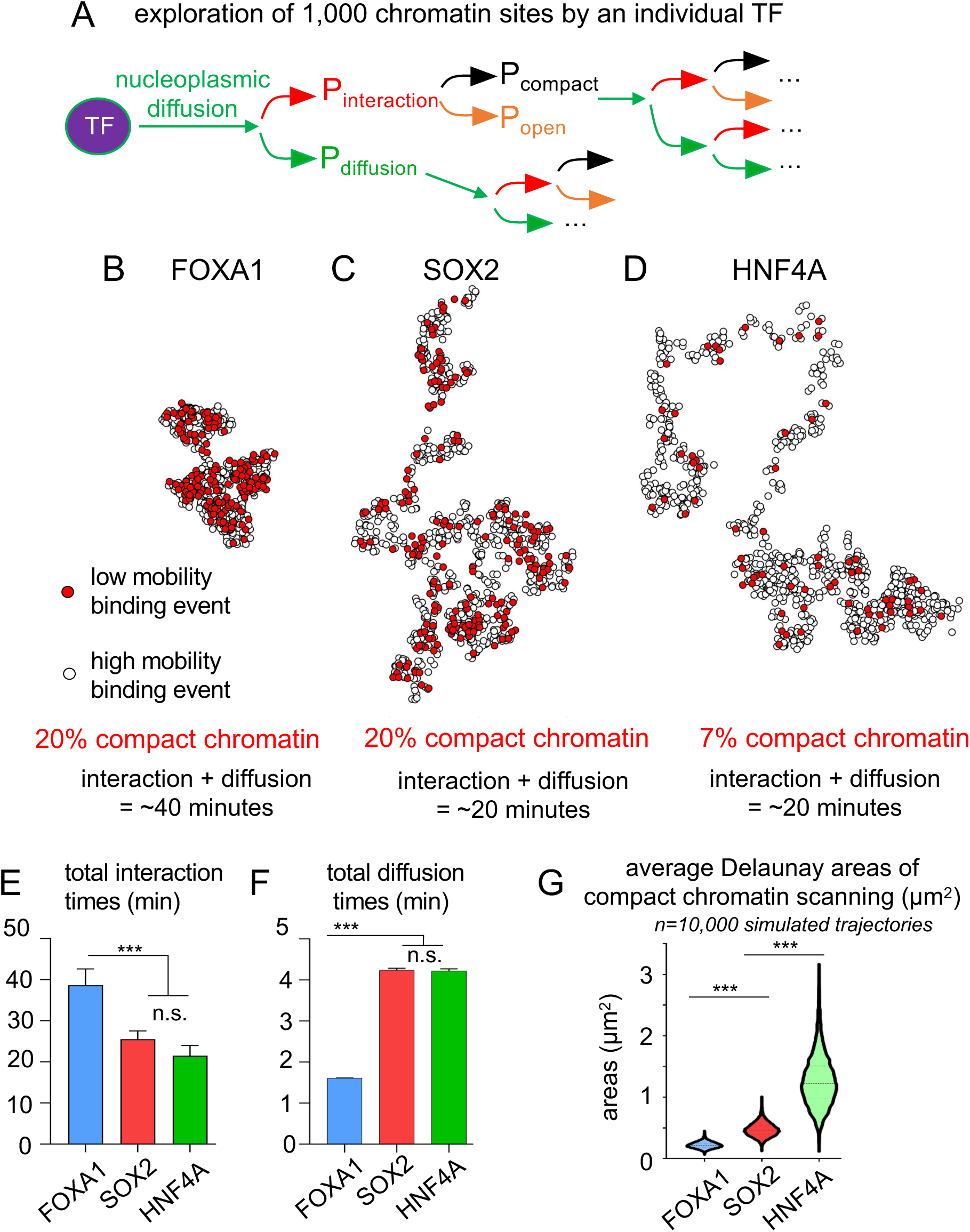
While displaying opposite kinetics, FOXA1 and SOX2 both perform a dense scanning of compact regions. A: Cartoon schematic of chromatin scanning trajectory simulations for 1,000 steps of a transcription factor (TF): after a step of nucleoplasmic diffusion (green), the probability that the TF diffuses again (P_diffusion_ green) or interacts with chromatin (P_interaction_, red) is inferred from the measurement of diffusion coefficients. If the TF interacts with chromatin, the probability to interact with a compact, low-mobility chromatin site (P_compact_, black) versus an open, high-mobility chromatin site (P_open_, orange) is inferred from the radius of confinement and average displacements measurements. The area scanned across the simulated trajectory is measured. B-D: Visualization of representative scanning trajectories by a single molecule of FOXA1 (B), SOX2 (C) and HNF4A (D) exploring 1,000 chromatin sites, using the algorithm in panel A. Each step of diffusion is set to occur in a random direction. Red dots indicate binding to low mobility, compact chromatin, while white dots indicate binding to high mobility, open chromatin. E: Total time spent interacting with chromatin during the exploration of 1,000 sites, inferred from the residence time distribution, for FOXA1, SOX2 and HNF4A. *** indicates p<0.0001, n.s. non-significant differences (p>0.05) as determined by one-way ANOVA, see Table S1). F: Total time spent diffusing in the nucleoplasm during the exploration of 1,000 sites, inferred from the average duration of diffusing tracks, for FOXA1, SOX2 and HNF4A. *** indicates p<0.0001, n.s. non-significant differences (p>0.05) as determined by one-way ANOVA, see Table S1). G: Measurement of average areas (µm^2^) after Delaunay triangulation of low-mobility, compact chromatin interactions spatial coordinates, for 10,000 simulated scanning trajectories of FOXA1, SOX2 and HNF4A. *** indicates p<0.0001, n.s. non-significant differences (p>0.05) as determined by one-way ANOVA, see Table S1).

Using our visualization tool, we find that FOXA1 (Figure 3B) elicits scanning of smaller territories than that of SOX2 (Figure 3C) and with slower temporal dynamics (Figure 3E and F). As indicated by similar P_diffusion_ and residence times, SOX2 and HNF4A present similar exploratory behaviors, but different capacities in engaging low-mobility chromatin and targeting silent domains (7% and 20%, respectively). To highlight how a 3-fold difference in the capacity to scan low-mobility, compact chromatin can result in an impaired targeting of silent domains, we inputted the rates of engagement with low-mobility chromatin in the trajectory for the three transcription factors (Figure 3B-D, red dots) and observed enhanced clustering of low-mobility compact chromatin scanning by SOX2, compared to that for HNF4A (Figure 3C-D, red dots).

To quantify such differences, we performed Delaunay Triangulation of compact chromatin interactions (Figure 3B-D, red dots and see STAR Methods) and used the areas of the Delaunay territories (Figure 3G and Supplemental Figure 5A and B) as a measure of the density in scanning of compact chromatin by transcription factors. Compared to HNF4A, FOXA1 and SOX2 presented a significantly denser scanning of low-mobility chromatin (Figure 3G), which could reflect how a critical density in low-mobility chromatin is necessary to achieve targeting of compact chromatin (see Discussion). The faster scanning behavior of SOX2 allows a larger exploration of chromatin domains, counterbalancing the difference in density seen between FOXA1 and SOX2, explaining how both pioneer factors present similar capacities in targeting silent chromatin.

Altogether, our simulations support how different modalities of engagement with low-mobility chromatin by FOXA1 and SOX2 result in site targeting in compact chromatin, as seen in ChIP-seq and CUT&RUN. As the probability of exploring the same site twice exists, in particular when driven by affinity to a consensus motif, the increased density of compact scanning could influence repetitive occupancy by pioneer factors in compact chromatin.

### Nonspecific DNA Interactions elicit slow chromatin scanning by FOXA1

We previously showed that the slow scanning behavior of FOXA1 was mainly governed by nonspecific DNA and nucleosome interactions ^17,40^. In order to assess whether perturbations of slow chromatin scanning by FOXA1 impairs the ability of the pioneer factor to interact with low-mobility, compact chromatin, we used previously characterized mutants ^17^ targeting amino acids within the DNA binding domain (DBD), responsible for specific (N216, H220 substituted with alanine, henceforth NHAA) or nonspecific (R262, R265 substituted with alanine, henceforth RRAA) DNA interactions (Figure 4A and Supplemental Figure 1A and 5C). The FOXA1-NHAA mutant has attenuated binding to a canonical FOXA motif, while FOXA1-RRAA mutants have attenuated binding to non-specific DNA sites, but can still bind specific sites, albeit weakly ^17,45^. Fast and SlowSMT of FOXA1-RRAA in human fibroblasts showed that loss of nonspecific DNA interactions causes a strong increase in diffusion (Supplemental Figure 5D and E) and a marked decrease in residence times (Supplemental Figure 5F and G). Conversely, loss of DNA site specificity (FOXA1-NHAA) had a lesser effect on diffusion and residence times (Supplemental Figure 5D-G), reflecting how nonspecific DNA interaction provide a major contribution to the slow chromatin scanning characteristic of FOXA1.

**Figure 4:**
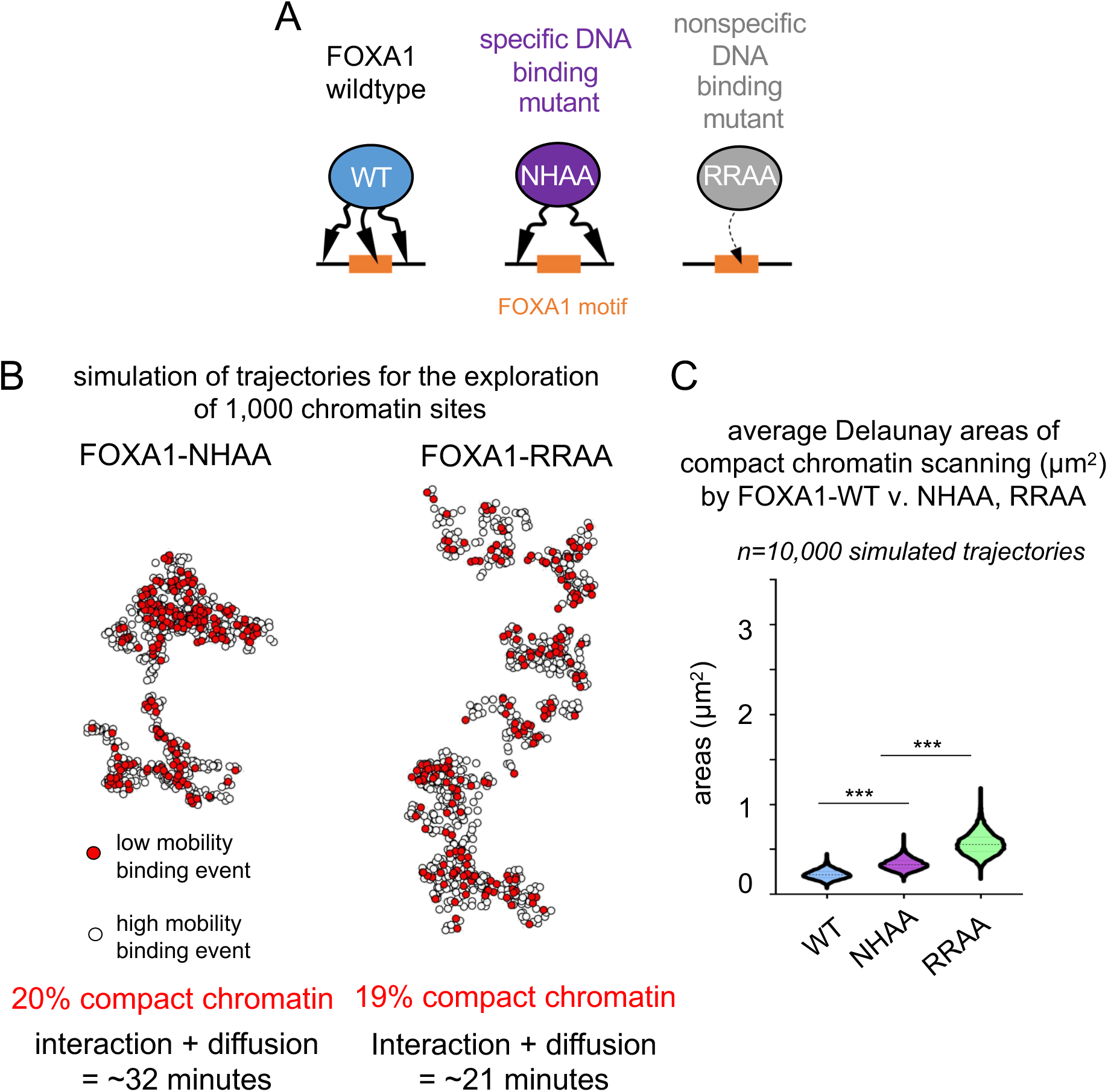
Impairing nonspecific DNA binding switches the scanning mode of FOXA1 from slow to fast without impairing interactions with low-mobility chromatin A: FOXA1-NHAA mutations target amino acids interacting with the DNA bases and abolish specific DNA binding. FOXA1-RRAA mutations target amino acids interacting with the DNA backbone and abolish nonspecific DNA binding B: Visualization of representative scanning trajectories by a single molecule of FOXA1-NHAA and FOXA1-RRAA C: Measurement of average areas (µm^2^) after Delaunay triangulation of low-mobility, compact chromatin interactions spatial coordinates, for 10,000 simulated scanning trajectories of FOXA1-WT, NHAA and RRAA. *** indicates p<0.0001, n.s. non-significant differences (p>0.05) as determined by one-way ANOVA, see Table S1).

Nevertheless, when measuring two-parameter mobility of the chromatin-bound mutants, we found that impairment of either nonspecific or specific DNA interactions had modest or no impact on scanning low-mobility chromatin (Supplemental Figure 6A and B). Thus, while impacting the dynamics of chromatin scanning by FOXA1, perturbation of specific or nonspecific DNA binding alone does not shape the extent of exploration of low-mobility, compact chromatin.

We simulated exploratory trajectories of FOXA1-NHAA and FOXA1-RRAA to illustrate how switches in exploratory behavior do not perturb the capacity of FOXA1 to target silent chromatin (Figure 4B). The loss of nonspecific DNA interactions (FOXA1-RRAA) results in an increased total size of explored areas, rates, and in a faster scanning of the chromatin sites (Figure 4B and Supplemental Figure 6C and D). Nevertheless, as FOXA1-RRAA conserved the high levels of engagement with low-mobility chromatin of FOXA1 (19%), the mutant achieved a dense chromatin scanning mode compatible with targeting of silent chromatin (Figure 4C) at levels similar to SOX2 (Figure 3C). Loss of specific DNA binding results in intermediate dynamics, closer to FOXA1-WT, confirming the major contribution of nonspecific DNA binding to the chromatin scanning by FOXA1 (Figure 4B and C, Supplemental Figure 6C and D). Altogether, our observations show that changing the slow exploratory mode of FOXA1 does not impair the capacity of the pioneer factor to scan low-mobility, compact chromatin.

### Pervasive, low-level sampling of most cognate motifs in the genome by FOXA1

A major question is whether pioneer factors recognize, or “sample,” many or most of their cognate motifs in any cell, despite forming clear binding peaks in particular cell types. Donaghey et al. ^11^ previously observed that FOXA1 weakly associates with sites stably bound among three alternative lineages, as assessed by a remnant ChIP-seq signal. Here, we assessed whether sampling by FOXA1 could be observed at all FoxA DNA motifs in the genome of human fibroblasts. We measured the input subtracted FOXA1 ChIP-seq signal at the 1,455,946 FoxA genomic motifs for which we did not identify a ChIP-seq peak (Figure 5A, B). We observed a weak but increased signal centered on FoxA motifs, suggesting that ectopically expressed FOXA1 samples many or most motifs that were not called as a peak (Figure 5A and B). Randomly-selected “background” sequences did not show specific enrichment for wild type FOXA1 or the mutants, confirming that the sampling occurs at motifs specific to the pioneer factor (Figure 5B, grey lines).

**Figure 5:**
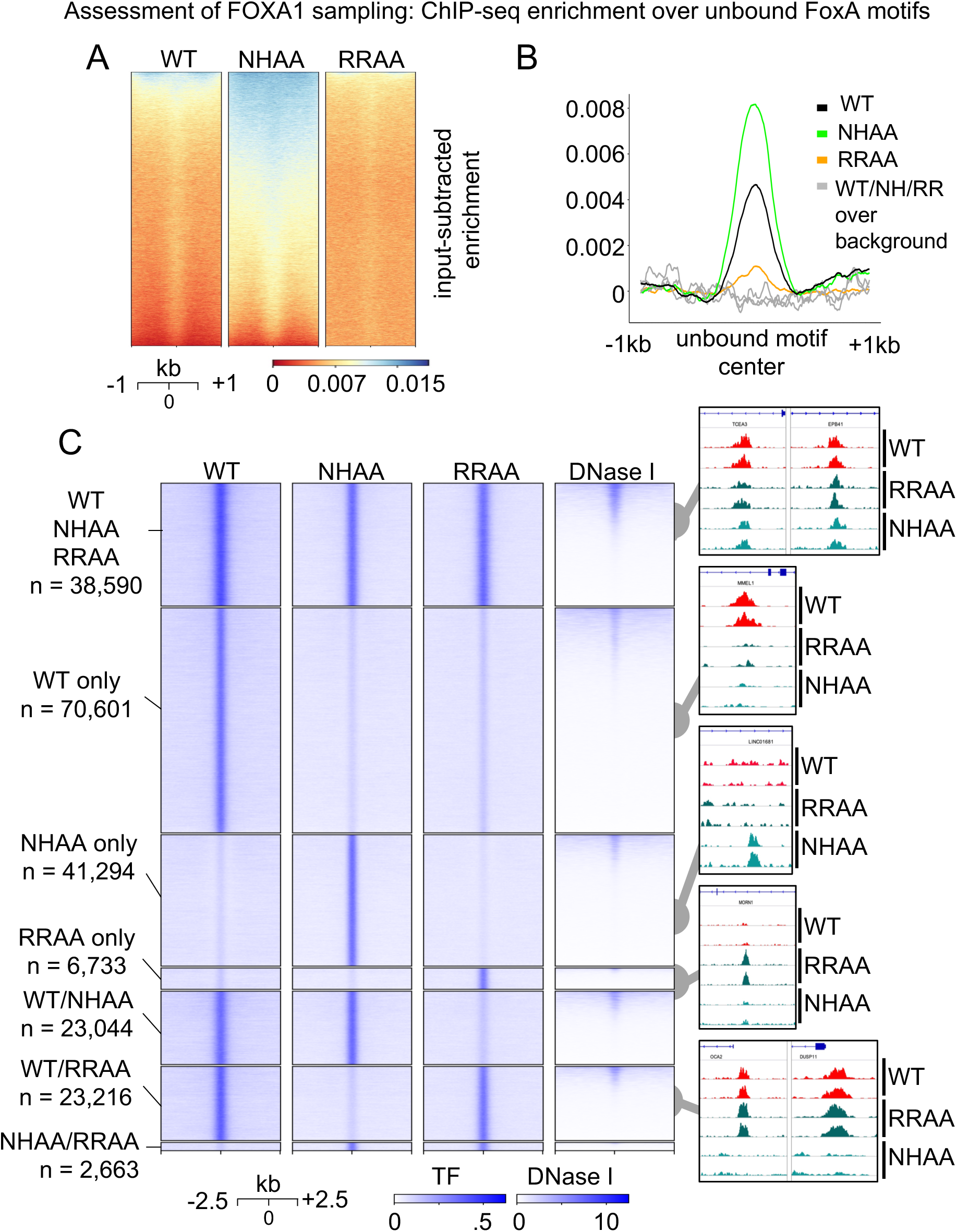
Impairment of specific and nonspecific DNA binding alters the motif sampling by FOXA1 but not targeting of silent chromatin. A: Heatmap of FOXA1-WT, FOXA1-NHAA, and FOXA1-RRAA ChIP-seq signal over unbound FoxA motifs. 1/10^th^ of all motifs (145,595) randomly sampled and plotted for visual clarity. B: Meta-analysis of ChIP-seq signal at all unbound FoxA motifs for FOXA1-WT (black), FOXA1-NHAA (green), FOXA1-RRAA (orange) compared to background sequences without a FoxA motif (gray). C: Heatmap displaying ChIP-seq and DNase I-seq signal and peak examples at sites bound by FOXA1-WT, FOXA1-NHAA and FOXA1-RRAA, alone or together.

To assess how changes in specific and non-specific DNA binding might impact site sampling by FOXA1, we performed ChIP-seq for FOXA1-NHAA-HALO and FOXA1-RRAA HALO in human fibroblasts (Figure 5A and Supplemental Figure 7A). As for wildtype FOXA1, we measured ChIP-seq levels at unbound FoxA motifs for FOXA1-NHAA and FOXA1-RRAA (Figure 5A and B). Loss of nonspecific DNA binding (RRAA) led to an attenuation of sampling by FOXA1 (Figure 5A heatmap and B, orange line, compare to WT black line). Impairing nonspecific DNA binding (RRAA) leads to a switch to a fast scanning mode by FOXA1, attenuating the sampling of FoxA motifs and relating to the fact that FOXA1-RRAA binds mainly to a subset of the wildtype chromatin sites (Supplemental Figure 7B). Unexpectedly, the specific DNA binding mutant (NHAA) displayed stronger signals over domains harboring a FOXA1 motif compared to wild type (Figure 5A and B, green line), reflecting an increase in sampling. Yet the increase in sampling signals extended widely from the FoxA motifs, consistent with a near absence of specificity and with the NHAA mutant targeting new specific and nonspecific chromatin sites (Supplemental Figure 7B), consistent with its slow scanning process (Figure 4B, C).

### FOXA1 targets compact chromatin via specific or nonspecific interactions

To understand how loss of specific versus nonspecific DNA binding impacts the targeting of wild type FOXA1 binding sites, we assessed the overlap of ChIP-seq peaks. We observed that loss of nonspecific or specific DNA binding by FOXA1 leads to targeting of a different, reduced set of sites (Figure 5C, Supplemental Figure 7B), as seen previously in NIH-3T3 cells ^46^. Still, 86% of FOXA1-RRAA targeted sites overlapped with FOXA1-WT and displayed a strengthened canonical FOXA1 motif compared to the wildtype peak set (Supplemental Figure 7C and D). Together with lower sampling levels (Figure 5B), impairment of nonspecific DNA binding leads to a reduced exploration of potential DNA binding sites. Conversely, only 58% of FOXA1-NHAA targeted sites overlap with FOXA1-WT and show a weakened enrichment for the FOXA1 motif, but a strengthening of other enhancer binding factor motifs, such as for AP-1 (Supplemental Figure 7C and E). As supported by the increased sampling (Figure 5B), Altogether, these results indicate that impairing specific or nonspecific DNA interactions impacts the search process for stable and transient DNA binding sites, which causes a redistribution of the FOXA1 cistrome.

We hypothesized that loss of specific or nonspecific DNA binding, by decreasing chromatin interactions, would also affect the capacity of FOXA1 to target silent, DNase-resistant chromatin. Nevertheless, we observed that the set of peaks bound by both FOXA1-NHAA and FOXA-RRAA (Supplemental Figure 7F) are largely in DNase I resistant chromatin sites, as for wild type FOXA1. Altogether with the conserved scanning capacity of low-mobility chromatin by FOXA1-NHAA and FOXA1-RRAA in SMT (Figure 4B and C, Supplemental Figure 6A and B), we conclude that the ability of FOXA1 to interact with compact chromatin occurs with either specific or nonspecific DNA interactions.

### Role of non-DBD domains in scanning and targeting by FOXA1 and SOX2

Given the defining characteristic of nucleosome binding for pioneer transcription factors, we evaluated whether the DNA binding domains of FOXA1 and SOX2, which are sufficient to bind nucleosomes *in vitro* ^47^, dominate the chromatin scanning and closed chromatin targeting characteristics of the full-length proteins.

We first assessed chromatin occupancy by the DBDs of FOXA1 and SOX2 (Figure 6A and B) by using CUT&RUN (SOX2-V5, Supplemental Figure 1A and 8A) or ChIP-seq (FOXA1-HALO, Supplemental Figure 1A and 8B). For both factors, the DBDs target a subset of the full-length sites, with the SOX2-DBD and FOXA1-DBD peaks binding 26% and 17% of full-length SOX2 and FOXA1 peaks, respectively (Figure 6C and Supplemental Figure 8C and D). Notably, both the SOX2-DBD and FOXA1-DBD showed a reduction primarily in targeting of DNase-resistant chromatin and generally targeted the more open chromatin sites seen by the full-length factors (Figure 6C and D, Supplemental Figure 8E and F).

**Figure 6:**
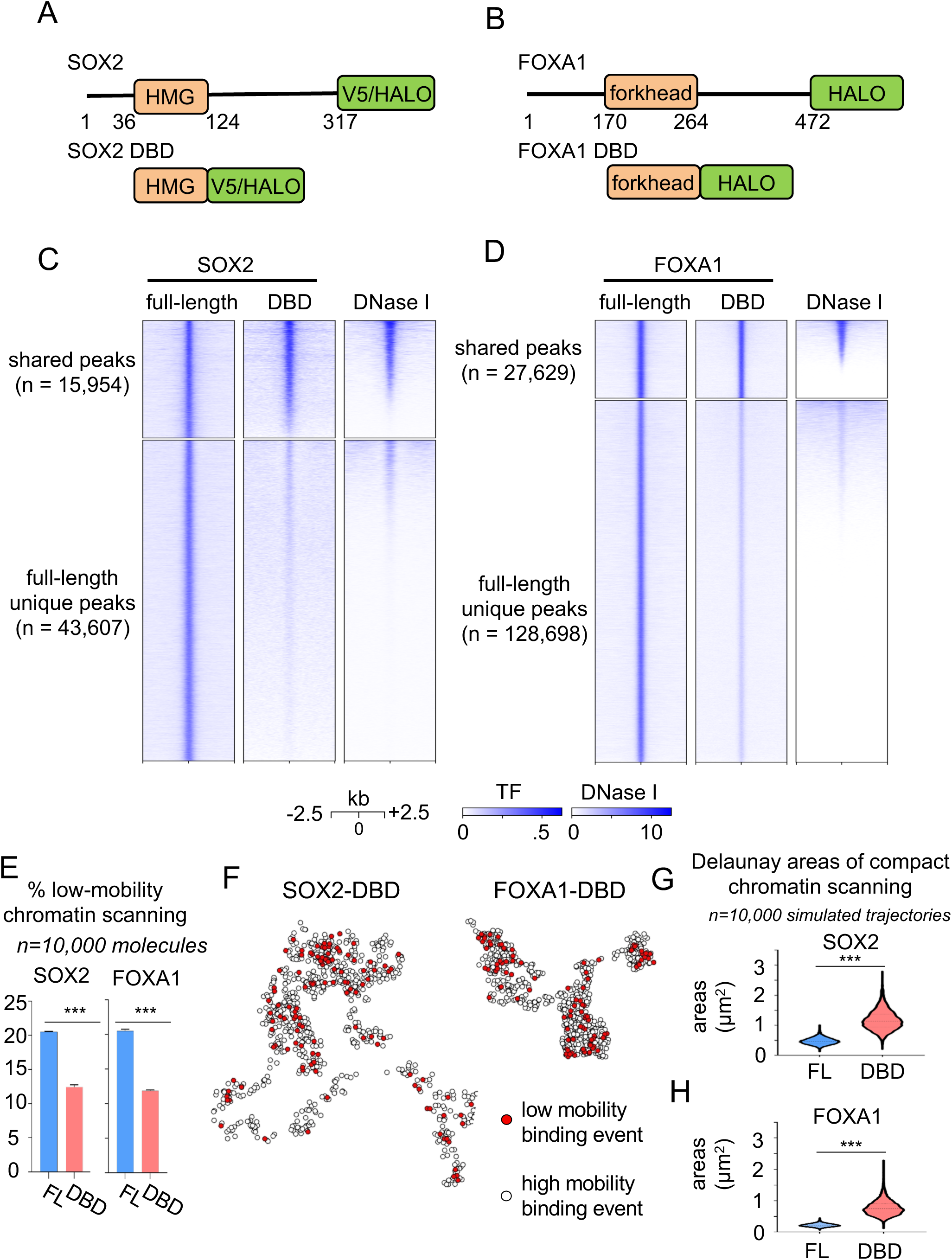
Non-DBD protein regions provide an essential contribution to the targeting and scanning of silent, compact chromatin by FOXA1 and SOX2. A-B: Graphic representation of SOX2 (A) and FOXA1 (B) -HALO/V5 constructs, and corresponding DBD truncations. C-D: Heatmaps displaying ChIP-seq and DNase I signals for SOX2, SOX2-DBD (C) and FOXA1, FOXA1-DBD (D). at shared peaks (upper panel) or peaks bound only by the full-length proteins. E: SMT measurement of scanning of low mobility, compact chromatin by HALOtagged SOX2 and FOXA1 full-length and DBD truncations. F: Visualization of representative scanning trajectories by a single molecule of SOX2-DBD and FOXA1-DBD. G-H: Measurement of average areas (µm^2^) after Delaunay triangulation of low-mobility, compact chromatin interactions spatial coordinates, for 10,000 simulated scanning trajectories of full-length or DBD truncations of SOX2 (G) and FOXA1 (H). *** indicates p<0.0001, n.s. non-significant differences (p>0.05) as determined by one-way ANOVA, see Table S1).

FastSMT diffusion coefficients and SlowSMT residence time measurements showed that the DBD domains alone exhibit impaired chromatin scanning (Supplemental Figure 9A). DBD molecules interact with chromatin (Supplemental Figure 9B and C). For SOX2 and FOXA1, the scanning activity of DBD truncations display a 35% (SOX2) and 55% (FOXA1) decrease in the levels of low-mobility chromatin scanning (Figure 6E, Supplemental figure 8D). Furthermore, truncations to the DBD resulted in a decrease of residence times, which was more marked for FOXA1 than SOX2 (Supplemental Figure 9E-G). While showing a lesser decrease in residence times than FOXA1, SOX2 still presented a strong, significant reduction in the number of stably-bound molecules (Supplemental Figure 9H).

Visualization of scanning trajectories displays how the loss of non-DBD domains results in the faster scanning of larger areas by FOXA1 and SOX2 (Figure 6F) and with a decrease in the density of low-mobility chromatin scanning (Figure 6G and H), correlating with the impaired targeting of silent chromatin states as seen by the genomic assays. Altogether, these results indicate that non-DBD domains provide a significant contribution to chromatin scanning, pioneering, and occupancy by FOXA1 and SOX2 by enhancing interactions with compact, low-mobility chromatin.

## DISCUSSION

In this study, we compared the chromatin binding and nuclear exploratory behavior of pioneer factors FOXA1 and SOX2 and non-pioneer factor HNF4A and discovered how distinct modes of chromatin scanning by the pioneer factors can lead to comparable targeting of sites in compact chromatin. FOXA1 and SOX2 primarily target DNase-resistant chromatin containing low-turnover nucleosomes, demonstrating that high nucleosome dynamics is not a pre-condition for pioneer factor binding. By performing SMT on the three transcription factors and using FOXA1 and SOX2 DNA binding domains alone, with attenuated targeting to DNase-resistant chromatin, we found that binding to silent chromatin targets involves interacting with sites that are spatially confined. Our visualization tool and simulation of pioneer factors scanning trajectories revealed how a key feature of pioneer factors for targeting silent sites is a critical density of interaction with low-mobility chromatin. Scanning high-density, compact scanning could enable repetitive occupancy by pioneer factors in the absence of a high residence time for each occupancy event. The faster scanning behavior of SOX2 might allow a larger exploration of chromatin domains, counterbalancing the difference in domain density scanning seen for FOXA1 and SOX2, and explaining how both pioneer factors present similar capacities in targeting silent chromatin.

Specifically, we found that FOXA1 and SOX2 reach their targets in compact chromatin with distinct scanning behaviors. FOXA1 is a slow explorer, showing more frequent and longer-lived interactions, while SOX2 is a fast explorer, diffusing frequently and interacting transiently with chromatin. The discrepancy between chromatin scanning kinetics of FOXA1 and SOX2 could relate to their mode of binding the nucleosome. While SOX2 recognizes an exposed, partial motif on the nucleosome surface ^10,13^, FOXA1 is predicted, via homology in its winged-helix domain, to compete with linker histone H1 ^26,47^, and to bind to nucleosomes in various orientations ^10^. The differential motif availability may cause SOX2 to move rapidly from nucleosome-to-nucleosome, while FOXA1 may be able to sample motifs in a mode that is less dependently of their position on the nucleosome. The MNase signal underlying DNase resistant peaks supports different modes of nucleosome binding: a strong MNase signal underlying SOX2 peaks suggests specific nucleosome orientations facilitate binding, while a fuzzy MNase signal underlying FOXA1 peaks suggests targeting nucleosomes that are not as well-positioned (Figure 1C-E). It will be interesting to determine which modalities are used by other pioneer factors and if various other modes of binding nucleosomes in vitro ^10^ predict in vivo chromatin targeting behavior.

We observed a discrepancy between the frequency of interactions with compact chromatin interactions measured by SMT (~20%) and the number of ChIP-seq or CUT&RUN peaks found in DNase I resistant chromatin (~70%). The observation highlights how chromatin interactions measured by FastSMT have a different nature, nonspecific and labile, of those assessed by genomics approaches, specific and stable. A corollary is that 80% of chromatin scanning interactions by FOXA1 and SOX2 seen by FastSMT occur in open chromatin, with only 30% of targeted open chromatin sites in ChIP-seq and CUT&RUN. This highlights the observed preference of FOXA1 and SOX2 for establishing stable interactions in compact chromatin, leading to the detection of a peak.

By various criteria, including nucleosome mapping, histone turnover, DNase resistance, and SMT, we observed markedly distinct, ectopic chromatin binding activities of FOXA1 and HNF4A in primary human fibroblasts, compared to another study using stably transfected, doxycycline-inducible constructs of FOXA1 and HNF4A in the tumor cell line K562 ^48,49^. K562 cells are multipotent, malignant, and aneuploid ^50,51^, possibly explaining differences observed and raising interesting questions about the nature of cancer cell chromatin. In our study, transcription factors were expressed de novo, typical of developmental or early reprogramming contexts in which FOXA1 functions with other factors to change cell fate ^52^.

We observed that the decrease of nonspecific DNA binding by FOXA1 leads to the loss of the slow chromatin exploration, but not of interactions with compact chromatin by ChIP-Seq. We observe that FOXA1-RRAA chromatin scanning behavior is very similar to SOX2, which is consistent with the fact that the mutation does not abolish FOXA1’s capacity to target silent chromatin, and that the slow scanning mode of FOXA1 might be imparted by other aspects of the pioneer transcription factor. The R262 of FOXA1 (the first R in the RRAA mutant) was recently found to be associated with prostatic cancer ^53,54^, highlighting how a switch in chromatin scanning dynamics could lead to the targeting of a new, pathogenic set of regulatory sites.

Given the sufficiency of the FOXA1 and SOX2 DBDs to bind nucleosome *in vitro* ^8^, we were surprised to find that both DBDs bind only a subset of the full-length sites, biased towards those in accessible chromatin. Correspondingly, SMT of the DBDs showed a shift in scanning low-mobility to high-mobility chromatin. The DBDs retain some ability to scan low-mobility chromatin and 30-40% of the bound sites are in DNase-resistant chromatin, showing that the DBD is capable of targeting nucleosomes in vivo, but is stabilized at many other sites by additional protein domains. An increasing literature identifies the role of non-DBD domains in stabilizing transcription factors within nuclear bodies and on chromatin ^55,56^. The difference in chromatin scanning modes between the full length and DBD-alone FOXA1 and SOX2 underlines the emerging appreciation that accession to distinct chromatin subtypes is facilitated by protein domains outside the DBD.

Numerous studies have individually assessed which chromatin states are bound by transcription factors ^11,12, 57–59^ and how the factors traverse the nucleus to reach their targets ^5,14,19,20,25,39,44^. Here, we profiled both chromatin targeting and single molecule trajectories to connect transcription factor mobility with binding of different chromatin states. We find that markedly different modalities of chromatin scanning lead to comparable extents of binding to compact chromatin with a low-mobility state. A further understanding of how pioneer transcription factors bind chromatin, beyond the level of the nucleosome, both through structural analyses as well as identifying how domains outside of the DBD interact with protein-partners and nuclear structures, will further shed light on the processes underlying the observed molecular behaviors and will enhance our ability to control cell fate at will.

## ACKNOWLEDGEMENTS

We thank Luke Lavis for kindly providing the Halo-ligand coupled to JF549 fluorophore. The research was supported by NIH grant R01GM36477 to K.S.Z.; NIH grant T32 GM008216 to A.K.

## AUTHOR CONTRIBUTIONS

Conceptualization: J.L, A.K, K.Z; Data Curation: J.L, A.K; Formal analysis: J.L, A.K; Funding acquisition: K.Z; Investigation: J.L, A.K, J.Z; Methodology : J.L, A.K; Project administration: K.Z, J.L; Resources: K.Z, J.L, J.Z; Software: J.L, A.K; Supervision: K.Z; Validation: J.L, A.K; Visualization: J.L., A.K, K.Z; Writing – Original Draft : J.L, A.K, K.Z

## DECLARATION OF INTERESTS

The authors have no conflicts of interest to declare.

## STAR METHODS

### RESOURCE AVAILABILITY

#### Lead Contact for Reagent and Resource Sharing

For other reagents generated in this study or questions about the reagents, please contact Ken Zaret.

#### Materials availability

All the materials generated in this study are accessible upon request.

#### Data and Code availability

The data and code used in this study are accessible upon request. All ChIP-seq, MNase-seq and CUT&RUN sequencing data has been deposited to the Gene Expression Omnibus (GEO) under the accession number GSE220570. DNase I-seq data was obtained from the ENCODE data portal (https://www.encodeproject.org/) with the following identifier: ENCSR000EME.

### EXPERIMENTAL MODEL AND SUBJECT DETAILS

#### Cell Lines and Tissue Culture

Human BJ fibroblasts cells were grown in High Glucose DMEM (ThermoFisher 11965118) pyruvate, GlutaMAX, supplemented with 10% characterized fetal bovine serum (Hyclone SH300071.03), 100 units/ml penicillin and 100 μg/ml streptomycin(Thermo Fischer 15140122) at 37C with 5% CO2.

Imaging experiments were carried out in Phenol red-free High Glucose Medium (ThermoFisher 21063029) pyruvate, GlutaMAX, in an imaging chamber heated at 37°C (more details in the Single Molecule Live Cell Imaging section).

For ChIP and CUT&RUN experiments, 1M of human BJ fibroblasts were seeded on four 15cm diameter plates. Once the cells attached, 5mL of non-concentrated rTTA2 lentivirus were added to each plate, in a total volume of 20mL of culture media supplemented with Polybrene (8ug/mL final concentration). The next day, the medium was replaced by 10mL of non-concentrated Halotag-TF virus, in a total volume of 20mL of culture media supplemented withPolybrene (8ug/mL final concentration) and doxycycline (1ug/mL final concentration). After 24h, the media was supplemented with fresh doxycycline at 1ug/mL final concentration.

### METHOD DETAILS

#### Plasmid Construction and Genome Editing

##### TETO-FUW plasmids (lentiviral vectors)

TETO-FUW-FOXA1-HALO/TETO-FUW-FOXA1-NHAA-HALO/TETO-FUW-FOXA1-RRAA-HALO/TETO-FUW-FOXA1-DBD-HALO/TETO-FUW-HNF4A-HALO/TETO-FUW-SOX2-HALO/TETO-FUW-SOX2-DBD-HALO/TETO-FUW-Histone H2B-HALO ORF of interest (see Key resource table) were PCR amplified with the adequate primers (see Table S1) and assembled using Gibson Assembly® Master Mix kit (NEB E2611L) with EcoRI (NEB) digested TETO-FUW-OCT4 (Addgene plasmid #20323).

#### Lentiviral production and concentration

Lentivirus were produced as described in ^60^. In brief, 293T cells were co-transfected with lentiviral expression vector, psPAX2 and PMDG. Fresh medium was added after 24h. After another 72h, the medium containing the lentivirus was centrifuges at 2,000 rpm for 10 min, passed through a 0.45 µm filter, pelleted by ultracentrifugation (24,000 rpm 3 hours) and resuspended at high concentration in 200 µL DMEM high glucose. Lentivirus were titered in H2.35 cells. Suboptimal M.O.I. (Multiplicity of Infection) was used (<1), in order to obtain low expression levels.

#### Western blotting

Nuclear extracts were performed as previously described ^61^, and run on 10 % Bis-Tris gels (Life technologies), followed by standard western blotting procedures. HALO and V5 tags were detected with a primary antibody (Promega G9211 1:1000 and Thermofisher R960-25 1:1000, respectively) and a anti mouse secondary antibody (Santa Cruz SC-2005, 1:10,000). Detection was performed with ECL Prime reagent (SuperSignal™ West Pico PLUS Chemiluminescent Substrate, ThermoFisher 34580) and the Amersham 600 imager.

#### MNase-seq

Profiling of compact nucleosomes was performed as previously described (Lim et al., 2023). Briefly, human fibroblasts were washed twice with PBS and dissociated with Trypsin. For each replicate, 200,000 cells were transferred to a 1.5mL tube and washed three times with wash buffer (200mM HEPES, 150mM NaCl, 0.5mM Spermidine, EDTA-free protease inhibitor). After the final wash, cells were permeabilized by resuspension in wash buffer with 0.05% digitonin for 5 minutes. Next, 80 U/mL MNase was added, and samples were left at 37C on a heating block for 2 minutes. The MNase reaction was activated by adding 3 mM CaCl2, and proceeded for 5 minutes at 37C. After 5 minutes, the digestion was halted by equal addition of 2x Stop Buffer (340 mM NaCl, 20 mM EDTA, 4 mM EGTA, 0.05% digitonin, 0.05 mg/mL RNase A, 0.05 mg/mL Glycogen). Digested chromatin was incubated for 30 minute at 37C to allow RNase activity, and then treated with 0.1% SDS and Proteinase K (200ug/mL) for an additional 2 hours. The resultant DNA was extract by phenol chloroform isolation. Verification of a nucleosome ladder was confirmed by running 250ng DNA on a 1.3% agarose gel.

Mononucleosome-sized DNA fragments were isolated by performing an AMPure XP bead selection. Sample volumes were adjusted to 50µl and mixed with 42.5µl AMPure XP beads (0.85x). After a 10 minute incubation at room temperature, followed by separation of the beads on a magnetic rack, the resultant supernatant was transferred to a fresh tube and the beads (containing larger DNA fragments) were discarded. The DNA was purified and concentrated by ethanol precipitation, resuspending the samples in 25µl TE. Isolation of the correct sized DNA fragments was confirmed by TapeStation analysis, and subsequently made into a library following manufacturers’ protocols.

#### MNase-seq analysis

Paired-end MNase-seq reads were trimmed with trim_galore version 0.4.3 with parameters --paired -q 20 --minimum-length 20. Trimmed reads were aligned to the human genome build hg19 using STAR aligner version 2.5.2a, with run parameters --alignSJDBoverhangMin 999 --alignIntronMax 1 --alignMatesGapMax 1000 --outFilterMultimapNmax 1. A quality-filtered BAM file was generated with the command samtools view -q 5 -f 2 -bS. BAM files were sorted by coordinate with command samtools sort. Bigwig files were generated using DANPOS3 command dpos.

#### CUT&RUN

CUT&RUN was performed as previously described, with minor adjustments (Skene et al., 2018, Janssens et al., 2018). Briefly, adherent fibroblasts were detached with Accutase, washed, bound to magnetic concanavalin A beads, and permeabilized with a dig-wash buffer (0.1% digitonin, 20mM HEPES pH 7.5, 75mM NaCl, 0.5mM Spermidine, EDTA-free protease inhibitor). Bead-bound cells were incubated with a V5 antibody (Thermo R960-25, 1:100) at 4C overnight. The following morning, the cells were washed twice with dig-wash buffer, incubated with pA/G-MNase for an hour at 4C, and washed twice more. After chilling cells on an ice block, MNase digestion was activated for 30 minutes in the presence of 2mM CaCl2. The reaction was stopped by adding an equal volume of 2X STOP buffer (340 mM NaCl, 20 mM EDTA, 4 mM EGTA, 0.1% Digitonin, 0.05 mg/mL glycogen, 5 mg/mL RNase A). Digested chromatin was extracted from permeabilized cells at 37C for 30 minutes, and DNA was purified by phenol-chloroform. Libraries were prepared as described (Liu et al., 2018).

#### CUT&RUN analysis

Paired-end reads were mapped to the human genome build hg19 using bowtie2 (v2.3.4.1) with run parameters --local --very-sensitive --no-unal --no-mixed --no-discordant -I 10 -X 700. A quality-filtered BAM file was generated with the command samtools view -q 5 -f 2 -bS. BAM files were sorted by coordinate with command samtools sort. Bigwig files for visualization were generated from BAM files with command bamCoverage --smoothLength 10 --normalizeUsing CPM --ignoreForNormalization chrM (deeptools 3.5.0). BED files were generated from BAMs using the bedtools command bamToBed (bedtools v2.27.1). BED files were converted to bedgraph format using the bedtools command genomecov. Peaks were called on the bedgraph files with SEACR v1.3 (Meers et al., 2019b), using the stringent setting and selecting the top 0.01% of regions by AUC. Final peak sets were selected by taking the union of biological replicates.

#### Chromatin immunoprecipitation

ChIP-seq was performed as in ^12^ with a total of 10M human BJ fibroblasts were used as a replicate for each ChIP-seq experiment. In brief, cells were washed twice with PBS before fixation in PBS-1% formaldehyde for 10 minutes. After quenching with 125mM Glycine, cells were washed three times in PBS and collected by scraping, and frozen at −80C. After three freeze-thaw cycles, the cell pellet was resuspended in 5mL of ice-cold hypotonic buffer (20 mM HEPES-KOH pH 7.5, 20 mM KCl, 1 mM EDTA, 10% Glycerol, 1 mM DTT, complete protease inhibitors cocktail) and incubated on a wheel for 10 min at 4C. After a 5 minute centrifugation at 2,000 rpm, the pellet was resuspended in 5 mL ice-cold lysis buffer (50 mM HEPES-KOH, pH7.5, 140 mM NaCl, 1mM EDTA, 10% glycerol, 0.5% NP-40, 0.25% Triton X-100, 1 mM DTT, complete protease inhibitors cocktail) and incubated on a wheel for 10 min at 4C. The cells were then dounced at least 5 times to isolate the nuclei and centrifugated 5 minutes at 2,000 rpm. The pellet was resuspended in 10 ml ice-cold wash buffer (Buffer III) (10 mM Tis-HCl, pH8, 200 mM NaCl, 1mM EDTA, 0.5 mM EGTA, 1 mM DTT, complete protease inhibitors cocktail) and incubated on a wheel for 10 min at 4C. After 5 minutes centrifugation at 2,000 rpm, the pellet of nuclei was frozen on dry ice, before thawing and resuspension in 2mL ice-cold sonication-lysis buffer (Buffer IV) (10 mM Tis-HCl, pH8, 100 mM NaCl, 1 mM EDTA, 0.5 mM EGTA, 0.1% Na-Deoxycholate, 0.5% N-lauroylsarcosine, 1 mM DTT, complete protease inhibitors cocktail). The nuclei were then sonicated 10 minutes on a COVARIS sonicator, leading t fragment sizes around 300 to 500 bp. Before the immunoprecipitation step, the chromatin extracts were solubilized with 1% triton and rocked at least 10 minutes, and the supernatant was saved after centrifugation at maximum speed for 20 minutes. For 1 replicate, the 2mL of chromatin from 10 million cells was immunoprecipitated with 12uL of anti Halotag antibody (Promega G9281) and rocked overnight at 4C. Before adding the antibody, 100uL total were uptake as inputs for each condition. The next day, 50 uL of dynabeads-protein G saturated overnight with 1mg/mL BSA in sonication buffer were added to the antibody/chromatin mix and rocked 3 hours at 4C. The beads were washed a total of 5 times with the ice-cold ChIP washing buffer (HEPES-KOH 50mM, LiCl 500mM, EDTA 1mM, NP-40 1%, NaDOC 0.7% + Complete EDTA-free from Roche), and once with Tris-EDTA. The beads and the inputs were then resuspended in 150 ul of ChIP Elution Buffer (1% SDS, 0.1M NaHCO3), and decrosslinked overnight at 65C. The next day, 150uL of Tris-EDTA was added to each tube, with 5uL of 10mg/mL RNase A and incubated 2 hours at 37C. 5uL of 10mg/mL Proteinase K were then added to the mix and incubated 1 hours at 55C. After a phenol chloroform extraction, ChIP DNA Clean and Concentrator columns (Zymo research) were used to concentrate the DNA in 30 uL of elution buffer. A total of 5 to 10ng of DNA were typically obtained, as measured by QUBIT and processed in libraries using NEBNext® Ultra™ II for DNA Library Prep (New England Biolabs).

#### ChIP-seq analysis

Single-end reads were aligned to the human genome build hg19 using bowtie2 v2.3.4.1 with run parameters --local -X 1000. A quality-filtered bam file was generated with the command samtools view -q 5 -bS (SAMtools v1.1). Optical and PCR duplicate reads were marked and removed using PICARD MarkDuplicates REMOVE_DUPLICATES=TRUE ASSUME_SORT_ORDER=queryname (GATK 4.2.6.0). Bigwig files were generated from BAM files with command bamCoverage --smoothLength 10 --normalizeUsing CPM --ignoreForNormalization chrM (deeptools 3.5.0).

BED files were generated from BAMs using the bedtools command bamToBed (bedtools v2.27.1). Peaks were called from BED files using MACS2 callpeak against a matched input control with a FDR of 0.1% (MACS2 v2.2.7.1). Peaks were then examined for overlaps with the ENCODE blacklist using bedtools intersect and were filtered accordingly. The union set of peaks from replicates were taken and used for downstream analyses.

H2B-HALO ChIP-seq experiments were spike-in normalized by aligning reads to a joint dm6/hg19 genome. The data was processed into BED files as described above. A scaling factor was calculated by dividing the number of uniquely-drosophila aligned reads by the number of uniquely-human aligned reads. Bigwig files were generated with the command bamCoverage --binSize 50 --scaleFactor ‘dm6/hg19 ratio’.

#### Motif analysis

Motif analysis was performed with HOMER-v4.6. Scanning for motif enrichment underlying FOXA1-HALO, HNF4A-HALo, or SOX2-V5 peak sets was performed using the command findMotifsGenome.pl with default parameters. Motifs differentially enriched in FOXA1-NHAA-HALO or FOXA1-RRAA-HALO mutants over wild type FOXA1-HALO was performed by setting the wild type peaks as the background set.

To explore sampling of FOXA1 we identified instances of the canonical FOXA1 motif across the genome by scanning for the position weight matrix FOXA1(Forkhead)/MCF7-FOXA1-ChIP-Seq(GSE26831)/Homer (Motif 110). FOXA1 motifs were then classified as unbound by selecting motif instances that do not overlap with the union set of FOXA1-NHAA/RRAA/WT peaks, using the command bedtools intersect -v.

#### Genomic data visualization

Heatmaps and metaplots were generated with deeptools version 3.5.0. Counts matrices were generated with command computeMatrix reference-point --missingDataAsZero --referencePoint center. Images were produced with the plotHeatmap or plotProfile commands, using default arguments.

#### Single Molecule Live Cell Imaging

All single-molecule live-cell imaging experiments were carried out in a Nanoimager S from Oxford Nanoimaging Limited (ONI), in a temperature and humidity controlled chamber, a scientific Complementary metal–oxide–semiconductor (sCMOS) camera with a 2.3 electrons rms read noise at standard scan, a 100X, 1.49 NA oil immersion objective and a 561 nm green laser. Images were acquired with the Nanoimager software.

30,000 human BJ fibroblasts were seeded in a LabTek-II chambered 8 well plates (Lab-Tek 155049) and infected with rTTA2 and the appropriate TETO-FUW-HALO lentivirus with 1 μg/ml doxycycline for 48h. The day of imaging, cells were treated with 5nM of Janelia Fluor 549 (JF549) HaloTag ligand (a kind gift from Luke Lavis, HHMI) for 15 minutes. Cells were subsequently washed three times in PBS at 37C, and Phenol Red-free High Glucose medium was added to each well. All imaging was carried out under HILO conditions (Tokunaga et al., 2008). For imaging experiments, one frame was acquired with 100ms of exposure time (10 Hz) to measure the intensity of fluorescence of the nuclei, and in Fast Single-Molecule Tracking (FastSMT) experiments, 5000 frames were acquired with an exposure of 10ms (100 Hz), while in SlowSMT, 200 frames were acquired with an exposure of 500ms (2 Hz).

### QUANTIFICATION AND STATISTICAL ANALYSIS

This protocol has been thoroughly described and explained in ^19^ and ^40^. All scripts are publicly available.

#### Two Parameters Single Molecule Tracking Analysis - Tracking algorithm

In brief, TIF stacks SMT movies were analyzed using MATLAB-based SLIMfast script (Teves et al., 2016), a modified version of MTT (Sergé et al., 2008), with a Maximal expected Diffusion Coefficient (DMax) of 3 μm2/s-1.

The SLIMfast output .txt files were reorganized by the homemade csv_converter.m MATLAB script (available in ^40^) in .csv format for further analysis.

#### Two Parameters Single Molecule Tracking Analysis - Classification of the tracks

The single molecule tracking .csv files (see previous section) were first classified by the homemade SMT_Motion_Classifier.m MATLAB script. Single molecule trajectories (or tracks) with a track duration shorter than 5 frames were discarded from the analysis. Motion tracks are classified by the script in different groups: tracks with α ≤ 0.7 were considered as Confined; motion tracks with 0.7 < α < 1 as Brownian; and motion tracks with α ≥ 1 as Directed. In addition, the motion tracks showing a behavior similar to a levy-flight (presenting mixed Confined and Directed/Brownian behavior) were detected by the presence of a jump superior to the average jump among the track + a jump threshold of 1.5, and classified as “Butterfly.” Butterfly motion tracks were segmented into their corresponding Confined and Directed/Brownian sub-trajectories for posterior analysis. As an additional filtering step of Confined motions (including confined segments of Butterfly tracks), we defined a jump threshold of 100nm, to filter out motion tracks with an average frame-to-frame jump size bigger than 100nm.

For the two-parameters analysis of all transcription factors, we defined the bound state as being the pool of Confined motion tracks and of the Confined segments of the Butterfly motion tracks. The unbound state was defined as the pool of Directed and Brownian motion tracks.

#### Two Parameters Single Molecule Tracking Analysis - Analysis of trajectories

After the track classification, the trajectories were analyzed as in ^19,40^ by the Two_Parameter_SMT.m homemade MATLAB script to quantify radius of confinement for the FastSMT bound states motion tracks only and average frame-to-frame displacement for FastSMT bound, unbound states motion tracks and SlowSMT motion tracks.

#### Two Parameters Single Molecule Tracking Analysis - Radius of Confinement versus Average displacement

For the joint representation, we have built scatter density plots using the same number of tracks for each condition (using random downsampling when necessary). For this purpose, we used the freely available Scatplot.m MATLAB function. We measured the percentage of particles in each previously defined chromatin mobility populations using the following gates:

- Very low mobility region: radius of confinement between 10 and 35 nm, average displacement between 10 and 29 nm.
- Low mobility region: radius of confinement between 35 and 50 nm, average displacement between 10 and 30 nm.
- Intermediate mobility region: radius of confinement between 10 and 35 nm, average displacement between 29 and 36 nm.
- High mobility region: radius of confinement between 35 and 55 nm, average displacement between 30 and 55 nm.
- Very High mobility region: radius of confinement between 55 and 300 nm, average displacement between 60 and 300 nm.

The fraction corresponding to compact chromatin scanning is the pool of very low and low mobility regions, preferentially bound by heterochromatin regulators (see ^19^.

#### Diffusion coefficients

The first 4 points of each T-MSD curve corresponding to each trajectory were fitted with a linear distribution to estimate the diffusion coefficient (Equation 2, Michalet, 2010):

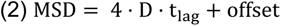

Where D is the diffusion coefficient, t_lag_ is the time between the two positions of the molecule used to calculate the displacement. The offset is due to the limited localization precision inherent to localization-based microscopy methods (~14 nm for our experiments). We set a coefficient of determination R^2^≥ 0.8 to ensure the good quality of the fitting performed to estimate D. Since the distribution of D follows a log-normal distribution ^63^, the Log_10_(D) was used for a proper visualization and fitting of the Gaussian Bi-modal distribution.

#### Residence times

We measured the residence times as performed previously ^64,65^. In brief, the “residence_time.m” Matlab script extracts the duration of every detected track and converted it in a residence time (Res.Time = Track_Duration·Exposure_Time).

The 1-cumulative distribution function (1-CDF) of the residence time of every detected track was fitted with a two-exponential decay equation on GraphPad Prism 8, to separate the 1-CDF in a short-lived and a long-lived population (Equation 3).

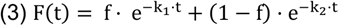

k_1_ and k_2_ are the unbinding constant rates in seconds^-1^, t_1_=1/k_1_ and t_2_=1/k_2_ the residence times in seconds, and f a number from 0 to 1 measuring the fraction belonging to each population. As photobleaching highly affects the measure of residence times, the measured k_1,2_ can be separated into their two contributions:

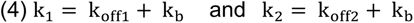

where k_off_ is the corrected unbinding rate and k_b_ the rate due to photobleaching. In order to measure k_b_, we used the k_off3_ of histone H2B, as in ^18,41^.

#### Visualization tool – generation of random walk coordinates

To build our visualization tool, we used a simplified version of the publicly available brownian_motion_simulation.m Matlab function from John Burkardt (function1.m, available here). This function generates a set of Euclidian coordinates from a position (0,0) using the following inputs: *t*, the time between each step of displacement; *m*, the spatial dimension; *s*, the step size in μm; and *n*, the number of steps.

The time step *t* is set as 0.01 seconds. The spatial dimension *m* is set to 2. To define the step size *s*, corresponding to the length of the diffusion distance between two interaction events, we measured the average displacement steps in the pool of unbound motion tracks (Directed + Brownian motion tracks, see section about track classification), by performing a lognormal fitting of the distribution (see Figure S4E-F). Here, we have used s=0.45 4m for all transcription factors.

To define *n*, we used the P_interacting_ (%) obtained by the measurement of Diffusion Coefficients.

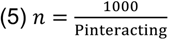

For example, for FOXA1, P_interacting_ =74%, thus *n*=1351 was used as input in function1.m, which thus provides a set of 1351 coordinates (output *ans*) with a step size of 0.45 μm. For SOX2, P_interacting_ =30%, thus *n*=3333 was used in the script.

The script viz_tool1.m uses the parameters defined in function1.m to randomly select 1,000 of the coordinates (output coord) generated by function 1, inputting the probability to interact with a chromatin sites versus performing another step of “free” nucleoplasmic diffusion at each step. The script viz_tool1 generates a visualization of the trajectory as a scatter plot.

#### Visualization tool – visualization of compact versus noncompact scanning

To visualize individual trajectories for transcription factors scanning 1,000 chromatin compact or open sites, we used the homemade viz_tool2.m script (available here). The script uses the set of coordinates (*ans*) from viz_tool1.m and its only input is Pcompact (%), corresponding to the percentage of compact chromatin scanning for each transcription factor. The viz_tool1.m script randomly select the corresponding percentage in the pool of 1,000 coordinates and display them as a red dot, representing a binding event to compact chromatin.

#### Visualization tool – simulation of chromatin scanning trajectories and measurement of compact scanning density

We used viz_tool3.m to generate 10,000 of the trajectories produced by viz_tool2.m, and measure the density of compact chromatin scanning, using Pcompact (%) as an input, similarly to viz_tool2.m.

To measure the density of compact chromatin scanning in the 10,000 trajectories simulated by viz_tool3.m, the script uses the Delaunay function of Matlab to triangulate the coordinates of compact chromatin scanning events (red dots in viz_tool2.m), and calculate the areas of the Delaunay territories (see Supplemental Figure 5A). The viz_tool3.m script then uses the publicly available fitExponential.m from Jing Chen to fit the distribution of Delaunay areas and obtain an average value for each trajectory (output “*fit”*), which were then plotted in violin plots using Graphpad Prism.

#### Visualization tool – time spent interacting with chromatin versus time spent free diffusing

The time spent interacting with chromatin was measured by randomly selecting 1,000 residence times from the distribution and summing them.

The time spent diffusing in the nucleoplasm was obtained by multiplying the input *n* of viz_tool1.m by the average duration of diffusing tracks (0.07 seconds, see Supplemental Figure 4G-H).

---

**Supplemental Figure 1 (related to Figure 1):**
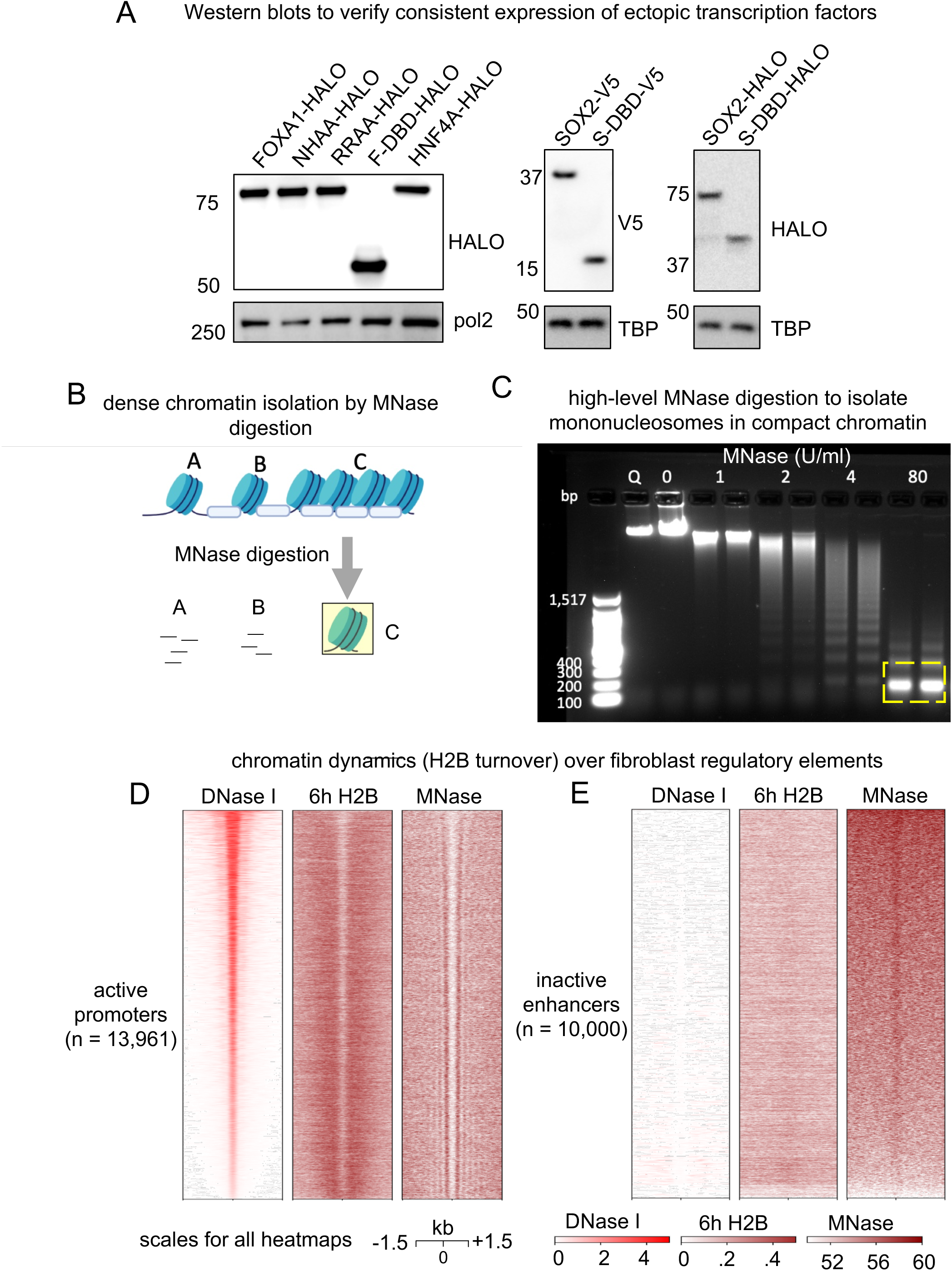
Expression of Transcription Factors and Histone H2B to assess Chromatin Targeting and Turnover. A: Western blot with an anti-HALO or V5 antibody, showing similar expression levels for FOXA1-HALO, FOXA1-NHAA-HALO, FOXA1-RRAA HALO, FOXA1-DBD-HALO, HNF4A-HALO, SOX2-V5, SOX2-DBD-V5, SOX2-HALO, SOX2-DBD-HALO, after 48 hours of doxycycline induction. Loading control : RNA polymerase II (noted pol 2). TATA-binding protein (noted TBP). B: Cartoon schematic of high-concentration MNase digestion to isolate mononucleosomes from compact chromatin regions. C: Chromatin digestion with varying MNase concentrations. Mononucleosome-sized DNA fragments marked by the yellow box were isolated for sequencing. D-E: Heatmaps displaying DNase I-seq, 6 hours H2B-HALO ChIP-seq and MNase-seq signals at active promoters (D) and enhancers inactive in human fibroblasts (E).

**Supplemental Figure 2 (related to Figure 1):**
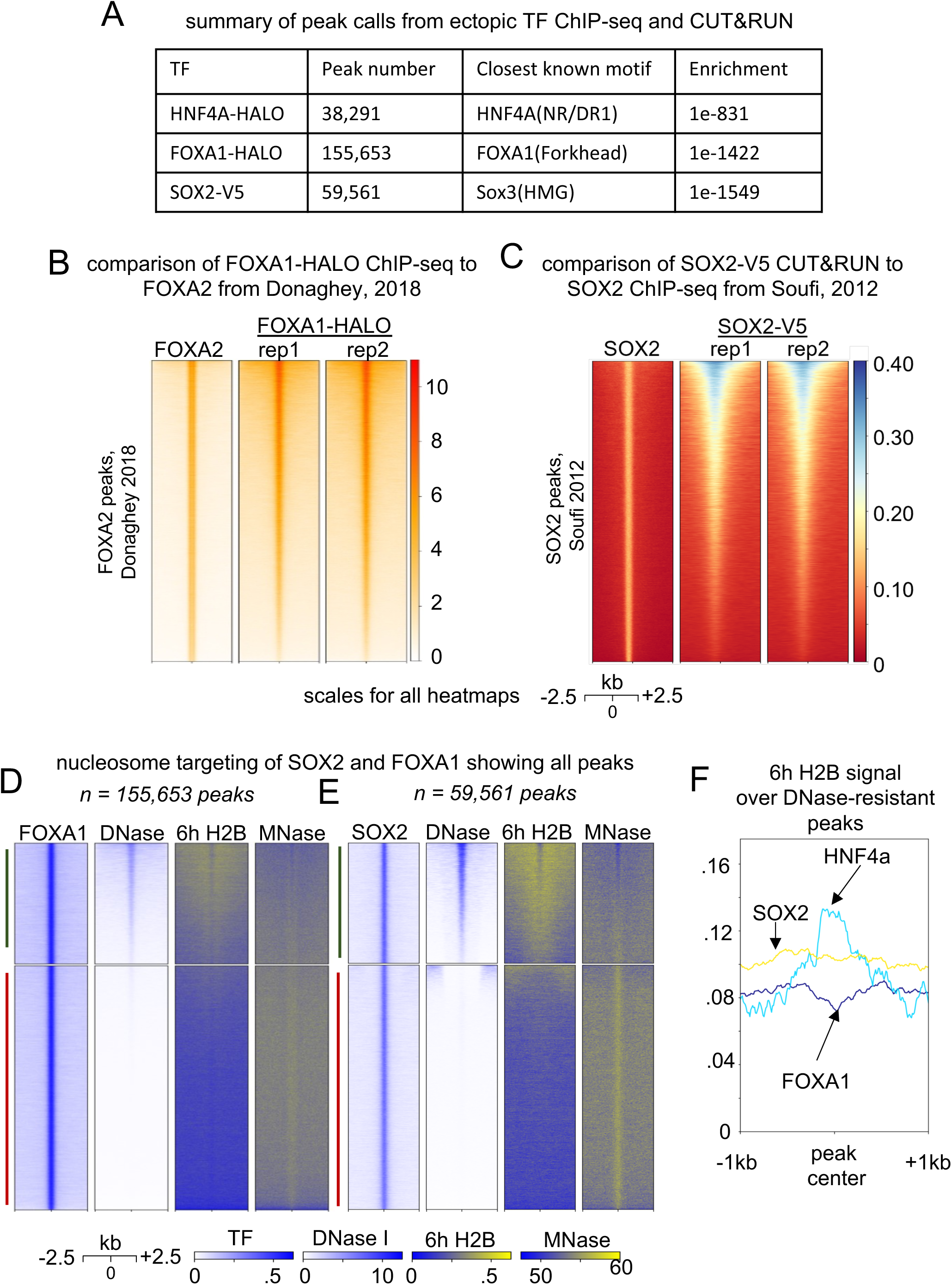
ChIP-seq of FOXA1, SOX2 and HNF4A in human fibroblasts. A: Number of peaks called, highest enriched motif and p-value after ChIP-seq of HNF4A-HALO, FOXA1-HALO and SOX2-V5. B-C: Comparison of ChIP-seq data from our experiments to previously published FOXA2 (B) and SOX2 (C) datasets. D-E: Heatmaps displaying FOXA1 ChIP-seq (D) and SOX2 CUT&RUN (E) signals at all peaks, with corresponding DNase I-seq, 6 hours Histone H2B-HALO ChIP-seq and MNase-seq signals. F: Meta-analysis of 6 hours Histone H2B ChIP-seq signal over peaks at DNase I-resistant sites for FOXA1, SOX2 and HNF4A

**Supplemental Figure 3 (related to Figure 2):**
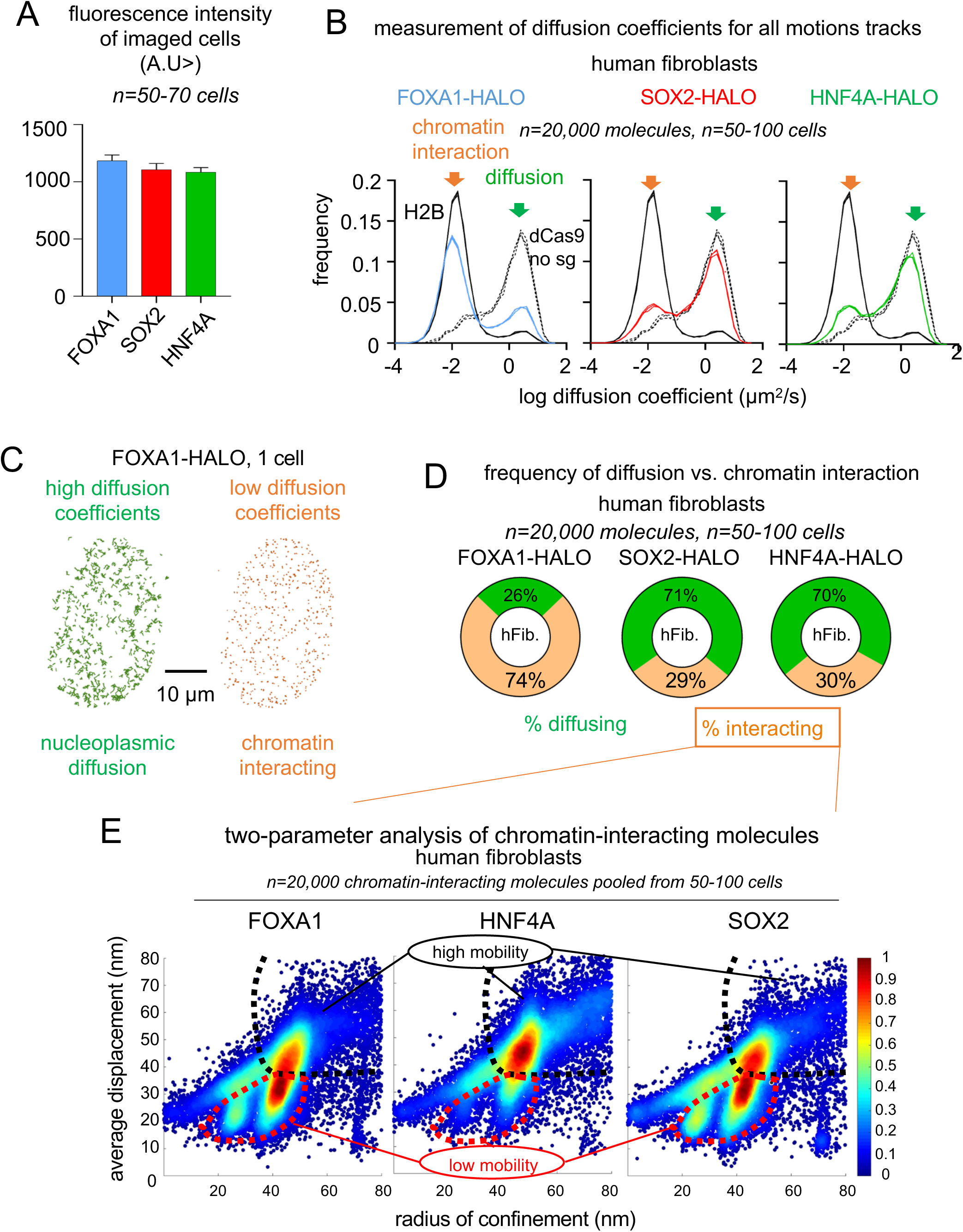
FastSMT of FOXA1-HALO, SOX2-HALO and HNF4A-HALO in Human Fibroblasts. A: Halo-549 fluorescence intensity (A.U.) of cells imaged for FOXA1-HALO, SOX2-HALO and HNF4A-HALO showing similar levels of expression. B: Logarithmic frequency of diffusion coefficients (μm^2^/s) of FOXA1-HALO (blue), SOX2-HALO (red) and HNF4A-HALO (green), in triplicates. In each panel, the logarithmic frequency of diffusion coefficients in triplicates for histone H2B and dCas9 expressed without a guide RNA is indicated with a hard or dotted black line, respectively Orange arrow: chromatin interacting molecules; Green arrow: molecules performing nucleoplasmic diffusion. n=20,000 molecules measured in 50-100 cells for each replicate. C: Aspects of high (green) and low (orange) diffusion coefficients motion tracks acquired over 50 seconds in a single nucleus. D: Frequency of nucleoplasmic diffusion and chromatin interactions of FOXA1-HALO, SOX2-HALO and HNF4A-HALO molecules, inferred from bimodal fitting of diffusion coefficient distributions of panel B. The values are the average of the triplicates. E: Scatter density plots of radius of confinement vs. average displacement for FOXA1-HALO, HNF4A-HALO and SOX2. The molecules interacting with low mobility, compact chromatin are encircled by a red dashed line, the molecules interacting with high mobility, open chromatin are on the right of the black dashed line.

**Supplemental Figure 4 (related to Figure 2):**
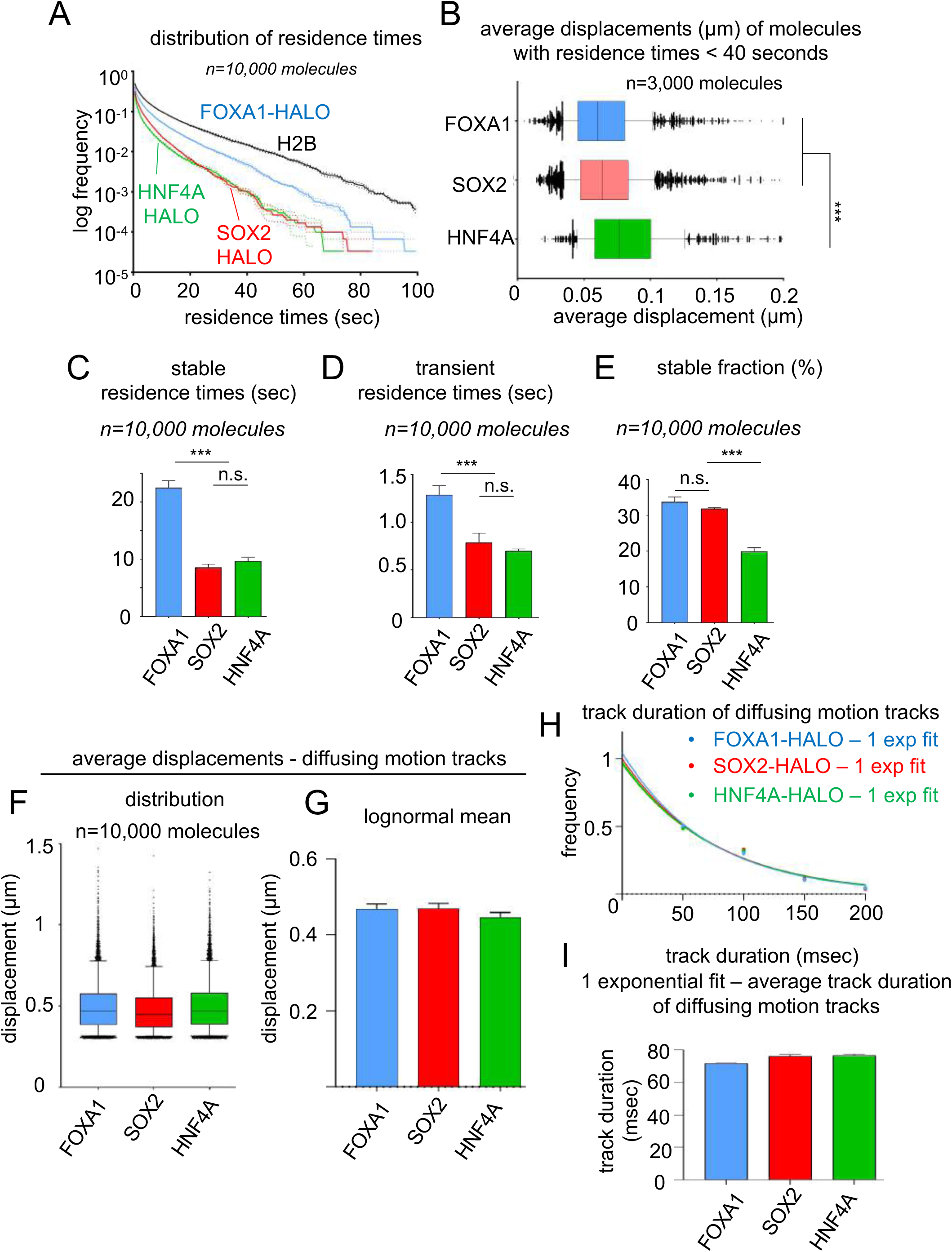
SlowSMT to measure residence times of FOXA1-HALO, SOX2-HALO and HNF4A-HALO. A: Logarithmic frequency distribution (1-CDF: cumulative distribution function subtracted to 1) of residence times for n=10,000 molecules of FOXA1-HALO (blue), SOX2-HALO (red) and HNF4A-HALO (green) and histone H2B (black), in triplicates. The hard line indicates the average frequency in each bin, and the dotted lines indicate the standard deviation. B: Average displacements (μm) of SlowSMT motion tracks of FOXA1 (blue) SOX2 (red) and HNF4A (green) with residence times below 40 seconds. C-E: 2-exponential decay fitting of the non-logarithmic residence time frequency Distribution provides for FOXA1-HALO, SOX2-HALO and HNF4A-HALO: average residence time (seconds) of the long (C) and short (D) - lived fraction, and size (%) of the long-lived fraction (E), in triplicates on n=10,000 molecules. Residence times values are corrected for photobleaching based on the residence times of Histone H2B. *** indicates p<0.0001, n.s. non-significant differences (p>0.05) as determined by one-way ANOVA, see Table S1). F-G: For FastSMT diffusing motion tracks of FOXA1-HALO, SOX2-HALO and HNF4A-HALO, distribution of average displacements (F) and means (G) after lognormal fitting of the distribution. H-I: For FastSMT diffusing motion tracks of FOXA1-HALO, SOX2-HALO and HNF4A-HALO, Logarithmic frequency distribution (1-CDF: cumulative distribution function subtracted to 1) of residence times, and 1-exponential decay fitting provides the average duration of nucleoplasmic diffusion events.

**Supplemental Figure 5 (related to Figure 3 and 4):**
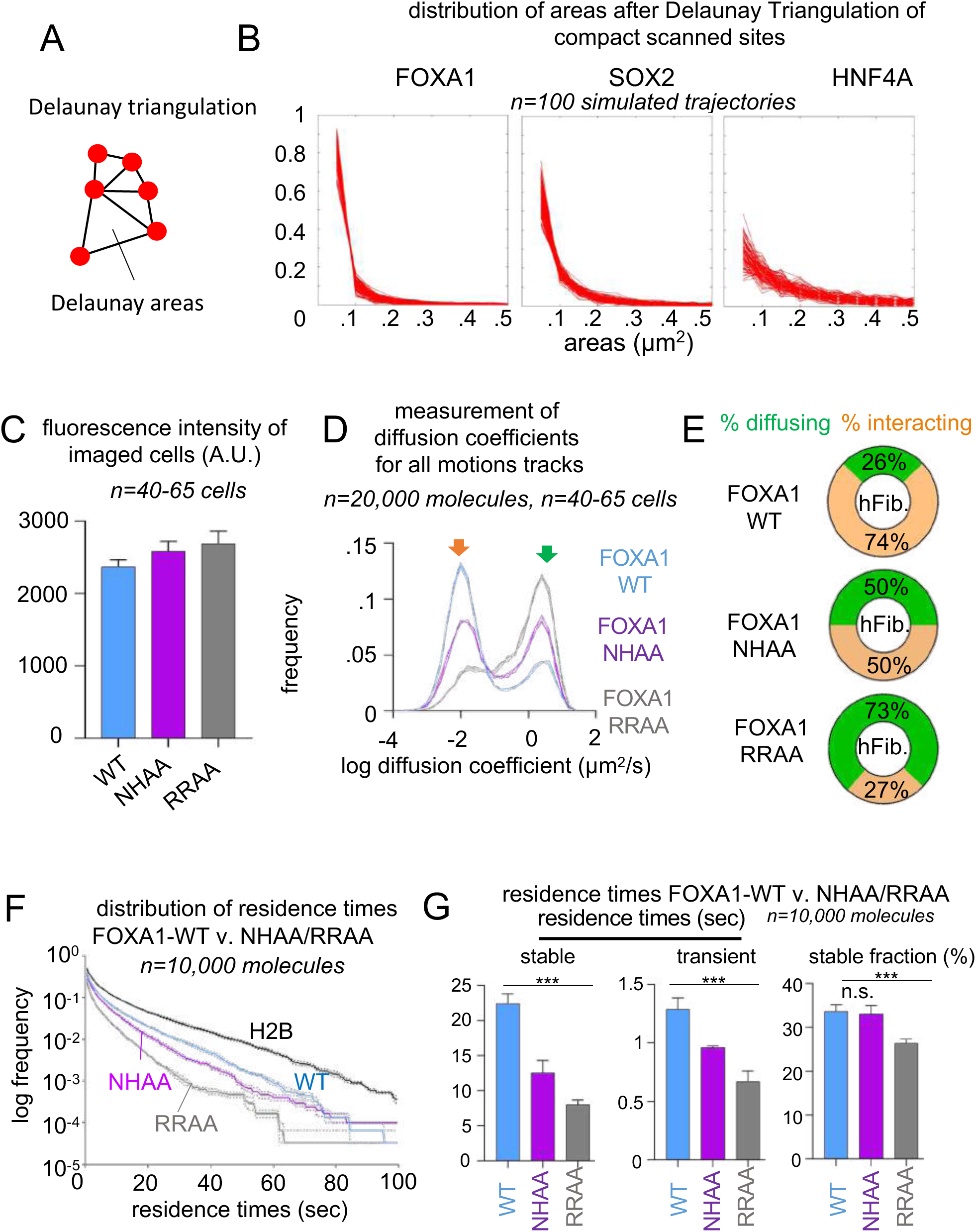
Measurement of SMT Parameters for FOXA1 DNA binding Mutants. A: Principle of Delaunay Triangulation B: Distribution of Delaunay areas after triangulation of low-mobility coordinates for 100 simulated trajectories. C: Halo-549 fluorescence intensity (A.U.) of cells imaged for FOXA1-HALO WT, NHAA and RRAA showing similar levels of expression. D: Logarithmic frequency of Diffusion Coefficients (μm^2^/s) of FOXA1-HALO-WT (blue), NHAA (purple) and RRAA (gray), in triplicates. Orange arrow: chromatin interacting molecules; Green arrow: molecules performing nucleoplasmic diffusion. n=20,000 molecules measured in 50-100 cells for each replicate. E: Frequency of nucleoplasmic diffusion and chromatin interactions of FOXA1-HALO-WT, NHAA and RRAA inferred from bimodal fitting of Diffusion Coefficient distributions of panel B. The values are the average of the triplicates. F: Logarithmic frequency distribution (1-CDF: cumulative distribution function subtracted to 1) of residence times for n=10,000 molecules of FOXA1-HALO-WT (blue), NHAA (purple), RRAA (grey) and histone H2B (black), in triplicates. The hard line indicates the average frequency in each bin, and the dotted lines indicate the standard deviation. G: 2-exponential decay fitting of the non-logarithmic residence time frequency Distribution provides for FOXA1-HALO-WT, NHAA and RRAA: average residence time (seconds) of the long and short-lived fraction, and size (%) of the long-lived fraction, in triplicates on n=10,000 molecules. Residence times values are corrected for photobleaching based on the residence times of Histone H2B.

**Supplemental Figure 6 (related to Figure 4):**
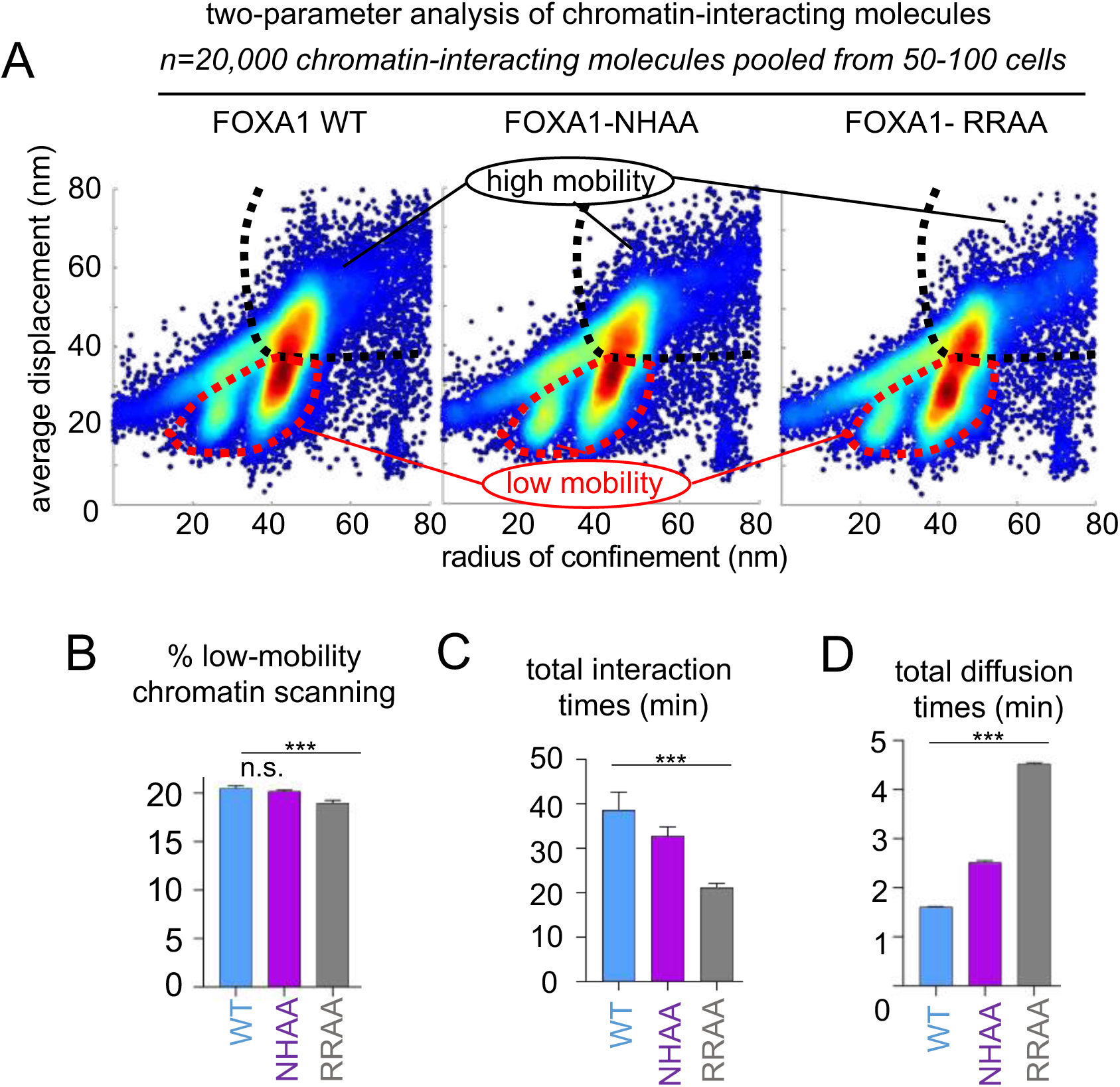
Low-Mobility Chromatin Interactions of FOXA1 DNA Binding Mutants. A: Scatter density plots of radius of confinement vs. average displacement for FOXA1-HALO-WT, NHAA and RRAA. The molecules interacting with low mobility, compact chromatin are encircled by a red dashed line, the molecules interacting with high mobility, open chromatin are on the right of the black dashed line. B: SMT measurement (%) of scanning of low mobility, compact chromatin by FOXA1-HALO-WT, NHAA and RRAA C: Total time (minutes) spent interacting with chromatin during the exploration of 1,000 sites, inferred from the residence time distribution, for FOXA1-HALO, -WT, NHAA and RRAA D: Total time (minutes) spent diffusing in the nucleoplasm during the exploration of 1,000 sites, inferred from the average duration of diffusing tracks, for FOXA1-HALO, -WT, NHAA and RRAA

**Supplemental Figure 7 (related to Figure 5):**
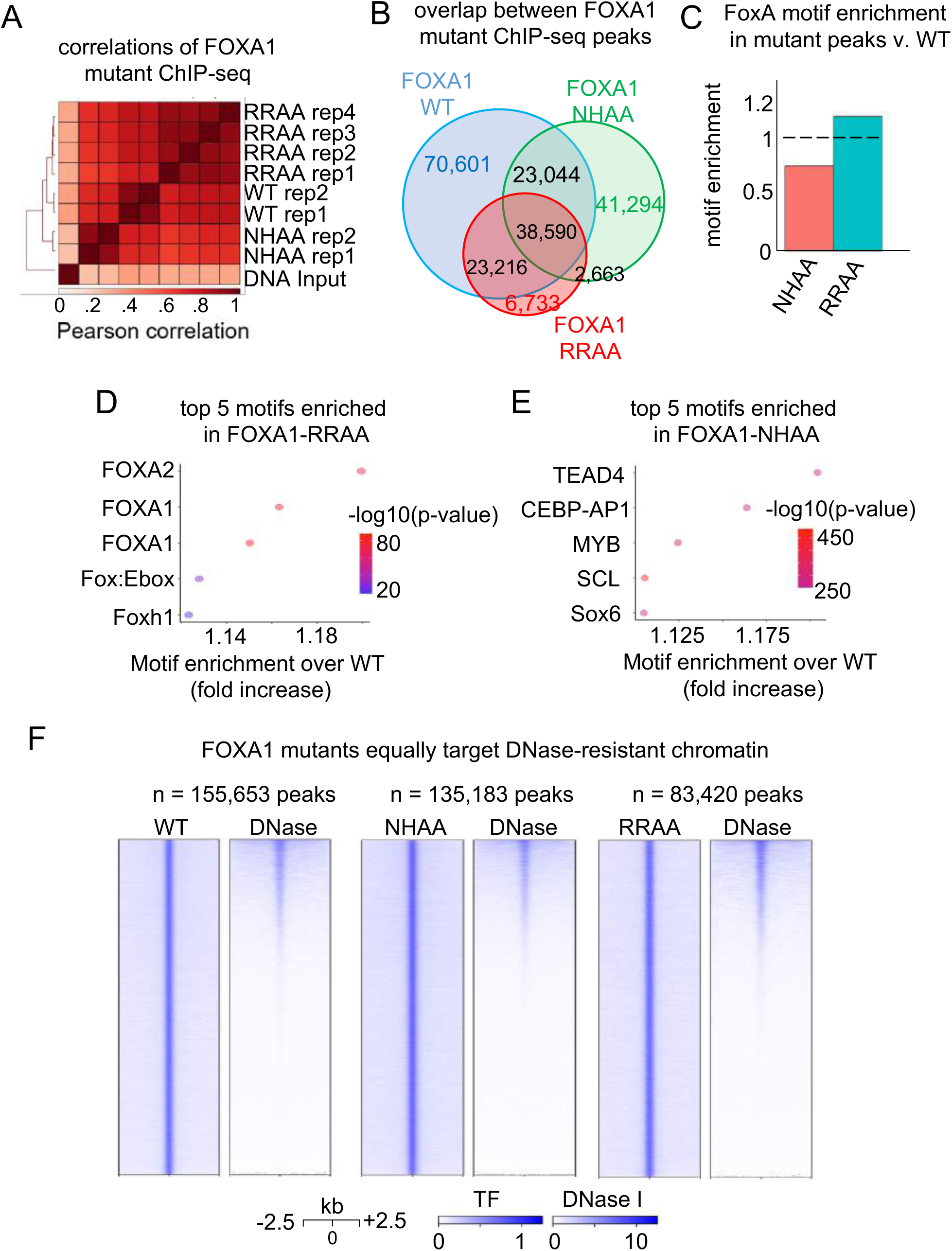
ChIP-seq of FOXA1 DNA Binding Mutants. A: Pearson Correlations of FOXA1-HALO-WT, NHAA and RRAA ChIP-seq replicates B: Venn diagram displaying overlapping between FOXA1-HALO-WT, NHAA and RRAA peak sets C: FOXA1 motif enrichment in NHAA and RRAA peaks versus FOXA1-HALO-WT D-E: Top 5 motifs found enriched in FOXA1-RRAA-HALO (D) and FOXA1-NHAA-HALO peak set. F: Heatmaps displaying HALO ChIP-seq signal and DNase I-seq signal at FOXA1-HALO-WT, NHAA and RRAA peaks.

**Supplemental Figure 8 (related to Figure 6):**
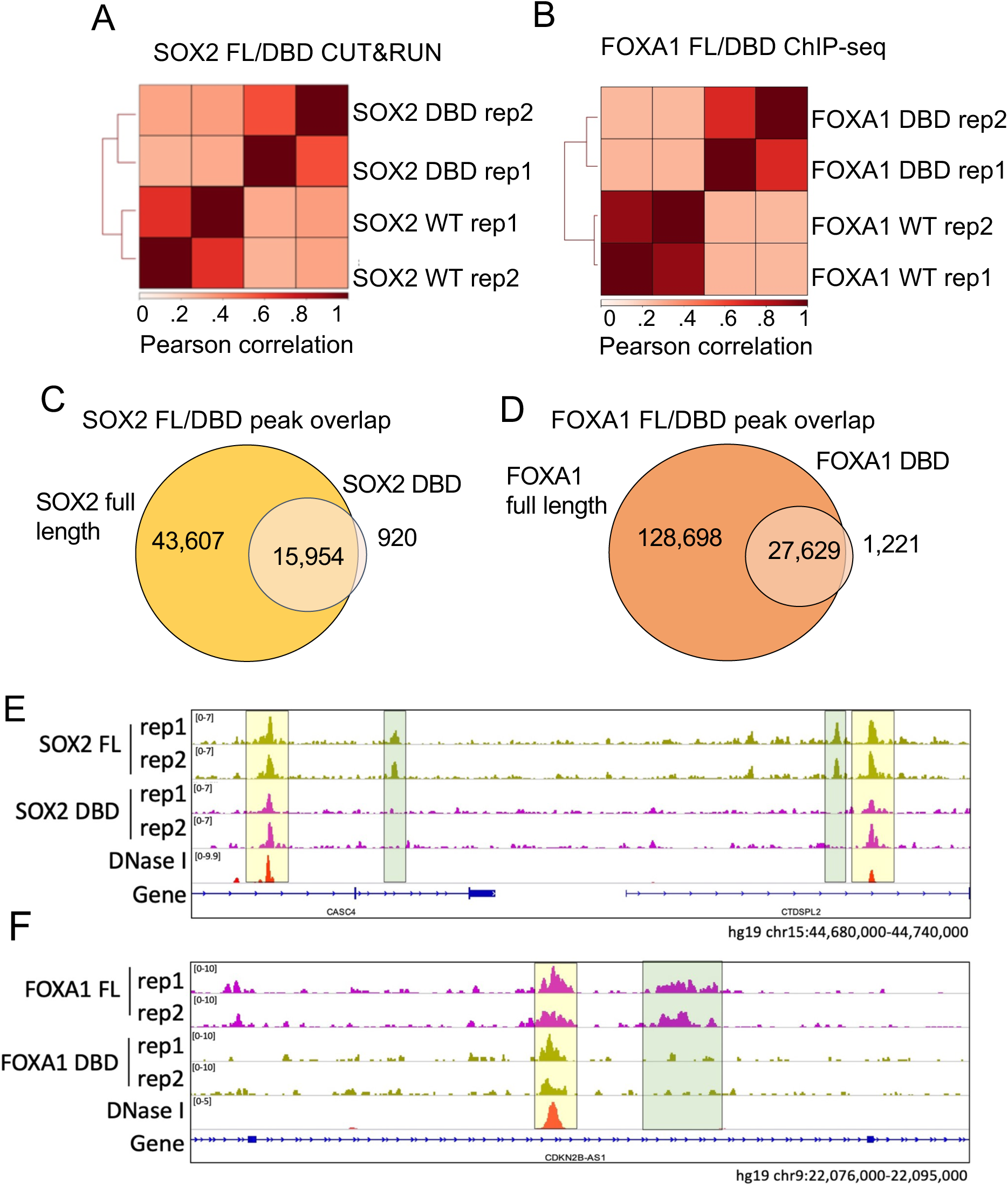
ChIP-seq of FOXA1 and SOX2 DNA Binding Domain Truncation. A-B: Pearson Correlations of SOX2-HALO-WT/DBD (A) and of FOXA1-HALO-WT/DBD (B) ChIP-seq replicates. C-D: Venn diagrams displaying overlapping between SOX2-HALO-WT and DBD (C) or FOXA1-HALO-WT and DBD (D) E-F: Representative example of sites bound in DNase I-resistant chromatin that the DBD truncation of SOX2 (E) and FOXA1 (F) do not bind to.

**Supplemental Figure 9 (related to Figure 6):**
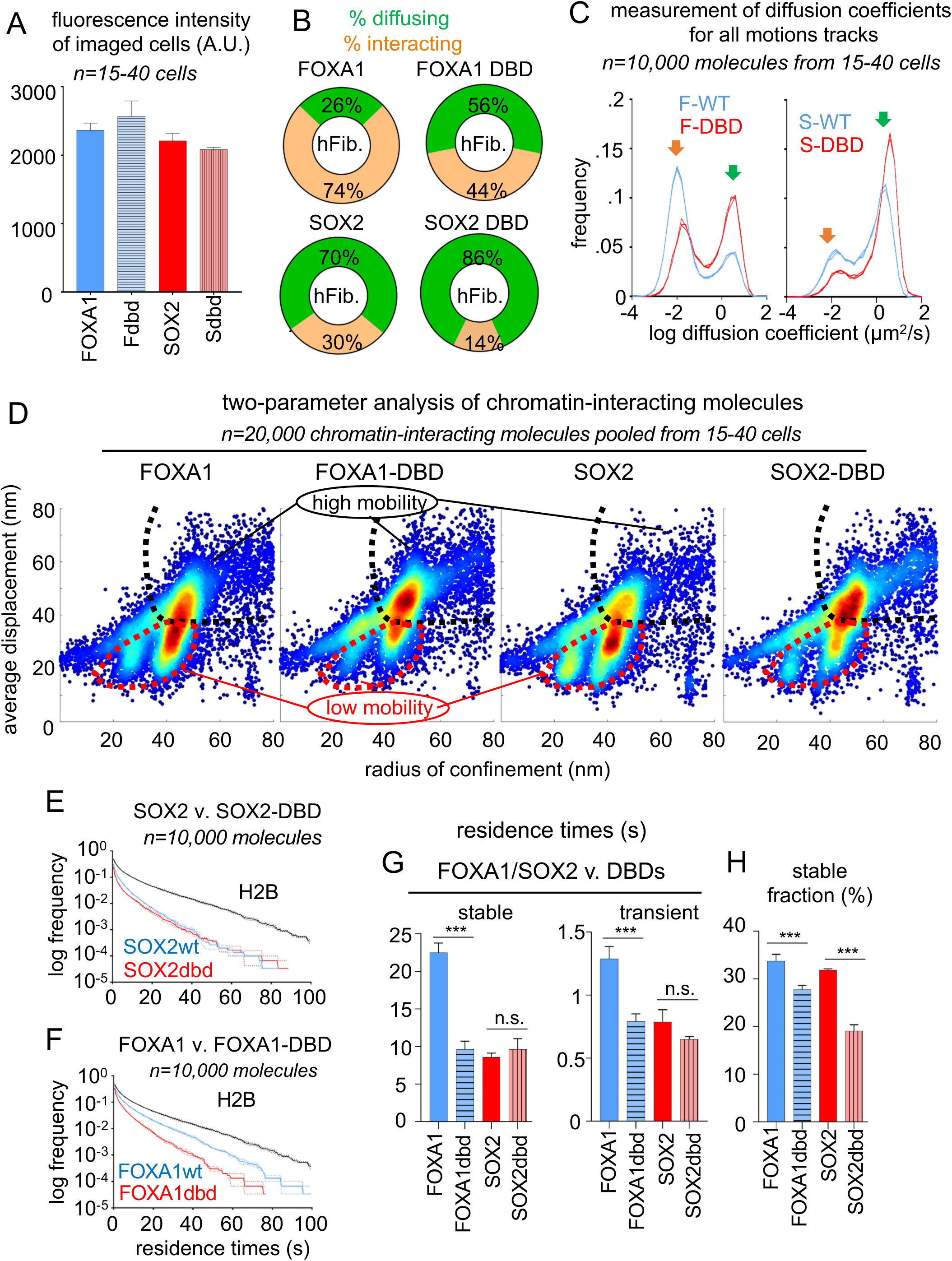
SMT of FOXA1 and SOX2 DNA Binding Domain Truncation. A: Halo-549 fluorescence intensity (A.U.) of cells imaged for FOXA1-HALO-WT/DBD and, SOX2-HALO-WT/DBD showing similar levels of expression. B: Logarithmic frequency of Diffusion Coefficients (μm^2^/s) of FOXA1-HALO-WT or SOX2-HALO-WT (blue) and DBD truncations (red) in triplicates. Orange arrow: chromatin interacting molecules; Green arrow: molecules performing nucleoplasmic diffusion. n=20,000 molecules measured in 50-100 cells for each replicate. C: Frequency of nucleoplasmic diffusion and chromatin interactions of FOXA1-HALO-WT, SOX2-HALO-WT and DBD truncations. The values are the average of the triplicates. D: Scatter density plots of radius of confinement vs. average displacement for FOXA1-HALO-WT, SOX2-HALO-WT and DBD truncations. The molecules interacting with low mobility, compact chromatin are encircled by a red dashed line, the molecules interacting with high mobility, open chromatin are on the right of the black dashed line. E-F: Logarithmic frequency distribution (1-CDF: cumulative distribution function subtracted to 1) of residence times for n=10,000 molecules of SOX2-HALO-WT (E) or FOXA1-HALO-WT (F) in blue or DBD truncations in red, in triplicates. The hard line indicates the average frequency in each bin, and the dotted lines indicate the standard deviation. G: 2-exponential decay fitting of the non-logarithmic residence time frequency Distribution provides for FOXA1-HALO-WT, SOX2-HALO-WT and DBD truncations: average residence time (seconds) of the long and short-lived fraction, and size (%) of the long-lived fraction, in triplicates on n=10,000 molecules. Residence times values are corrected for photobleaching based on the residence times of Histone H2B. *** indicates p<0.0001, n.s. non-significant differences (p>0.05) as determined by one-way ANOVA, see Table S1).

## Notes

### Competing Interest Statement

The authors have declared no competing interest.

### Summary of Updates

Formatting alteration, no changes to results presented.

